# Exploring the Interplay between BOLD Signal Variability, Complexity, and Static and Dynamic Functional Brain Network Features during Movie Viewing

**DOI:** 10.1101/2023.02.28.530546

**Authors:** Amir Hossein Ghaderi, Hongye Wang, Andrea Protzner

## Abstract

As the brain is dynamic and complex, knowledge of brain signal variability and complexity is crucial in our understanding of brain function. Recent resting-fMRI studies revealed links between BOLD signal variability or complexity with static/dynamics features of functional brain networks (FBN). However, no study has examined the relationships between these brain metrics. The association between brain signal variability and complexity is still understudied. Here we investigated the association between movie naturalistic-fMRI BOLD signal variability/complexity and static/dynamic FBN features using graph theory analysis. We found that variability positively correlated with fine-scale complexity but negatively correlated with coarse-scale complexity. Hence, variability and coarse-scale complexity correlated with static FC oppositely. Specifically, regions with high centrality and clustering coefficient were related to less variable but more complex signal. Similar relationship persisted for dynamic FBN, but the associations with certain aspects of regional centrality dynamics became insignificant. Our findings demonstrate that the relationship between BOLD signal variability, static/dynamic FBN with BOLD signal complexity depends on the temporal scale of signal complexity. Additionally, altered correlation between variability and complexity with dynamic FBN features may indicate the complex, time-varying feature of FBN and reflect how BOLD signal variability and complexity co-evolve with dynamic FBN over time.

## 1 Introduction

The human brain is a complex dynamic system, the state of which depends on both current and past experiences. That is, the brain generates transient states that are metastable, and represent the brain’s dynamic repertoire of functional configurations. Possible configurations are based on anatomical structure, current internal or external input, and past experience (Deco, Jirsa, & McIntosh, 2011; Rabinovich et al., 2012). The brain is also complex in that it embodies a repertoire of many networks which jointly segregate and integrate incoming information to co-ordinate behaviour (Tononi, Sporns, & Edelman, 1994). Because brain activity is time-varying and complex, exploring the dynamics and complexity of brain signal is critical to advancing our understanding of brain function.

To characterize the dynamic features of brain signal, some functional magnetic imaging (fMRI) work has shifted focus from examining measures of central tendency (e.g., mean activity) to examining measures of variability. The most commonly used metric is the standard deviation (SD) of blood oxygen level dependent (BOLD) signal (BOLD SD; Garrett, Kovacevic, McIntosh, & Grady, 2010, 2011), which reflects transient, within-individual temporal fluctuation of the signal. Another metric, multiscale entropy (MSE; Costa, Goldberger, & Peng, 2002, 2005), is more commonly used with EEG and focuses on entropy-based complexity measurement (i.e., brain signal complexity). MSE measures the predictability or repetitive structure in a time series, across multiple, coarse-grained time scales. Previous work suggests that MSE at fine time scales is related to local brain information processing, whereas MSE at coarse time scales is associated with long distance communication between distributed brain regions (McDonough & Nashiro, 2014; McIntosh et al., 2014).

Theoretical and empirical work has shown that brain signal variability and complexity are crucial for information processing and brain functional network reconfiguration (Deco, Jirsa, McIntosh, Sporns, & Kötter, 2009; Deco et al., 2011; Deco & Jirsa, 2012; Garrett et al., 2010, 2011; Ghosh, Rho, McIntosh, Kötter, & Jirsa, 2008; McIntosh, Kovacevic, & Itier, 2008; McIntosh et al., 2014; Protzner, Kovacevic, Cohn, & McAndrews, 2013). An optimal amount of variability or complexity may facilitate exploration of different interaction routes for separate regions and allow for switching between available functional networks spontaneously. These processes help the brain to quickly settle into the best functional network configuration to respond to any given input (Deco et al., 2009, 2011, 2012; McIntosh et al., 2010). Although both brain signal variability and complexity reflect brain dynamics, it is worth noting that variability and complexity are not equivalent (Easson & McIntosh, 2019; Lipsitz, 2002; Van Emmerik, Ducharme, Amado, & Hamill, 2016). Complex signal demonstrates variability, but variable signal may not be complex (https://archive.physionet.org/tutorials/cv/).

Empirical work examining resting state signal has linked both brain signal variability and complexity to structural and functional connectivity (Baracchini et al., 2021; Easson & McIntosh, 2019; McDonough & Nashiro, 2014). For example, a resting state fMRI study examined how BOLD signal variability and sample entropy link to global efficiency (GE), which is a graph theory metric that represents the overall information processing capacity of structural brain connectome (Easson & McIntosh, 2019). They identified a positive correlation between all three measures, suggesting that information processing capability of structure and functional networks are related. More recent fMRI work also showed that when brain region pairs show stronger functional connectivity, BOLD signal variability in the corresponding regions was more similar in value regardless of anatomical distance and structural difference between the two regions (Baracchini et al., 2021). Finally, using MSE, McDonough and Nashiro (2014) demonstrated that the relationship between complexity and functional connectivity is scale dependent. They identified a negative correlation at fine scales (approximately from 1 to 2.8 second windows) but a positive correlation at coarse scales (approximately from 4.9 to 18 second windows; McDonough & Nashiro, 2014). The authors related this finding to a neural model of information processing (Baptista & Kurths, 2008), which suggested that information transmission capability of a channel (comparable to information transmission capability as indexed by MSE) is negatively associated with neuronal phase synchronization (comparable to functional connectivity) at fine time scales and is positively associated with phase synchronization at coarse time scale.

The abovementioned studies used static functional connectivity measures, which represent average functional connectivity over relatively large time windows (e.g., 5-10 minutes). More recent work suggests that functional connectivity is dynamic, and reconfigures over time on the order of milliseconds to minutes (Allen et al., 2014; Battaglia et al., 2020; Chang & Glover, 2010; Di & Biswal, 2015; Handwerker, Roopchansingh, Gonzalez-Castillo, & Bandettini, 2012; Hutchison, Womelsdorf, Gati, Everling, & Menon, 2013; Kang et al., 2011; Kiviniemi et al., 2011; Van de Ville, Britz, & Michel, 2010). Importantly, dynamic functional connectivity analyses disclose transient interplay between brain networks, which is overlooked when using the more traditional static functional connectivity measures (Hutchison et al., 2013). Since brain signal variability, complexity and *dynamic* functional connectivity all reflect time-varying features of the brain signal, it is likely that the relationship between them is stronger or different than those between brain signal variability, complexity and *static* functional connectivity. Additionally, the relationship between variability, complexity, and *dynamic* functional connectivity may better characterize brain dynamics and transient functional states. In fact, the complex reconfiguration of dynamic functional connectivity during resting state is associated with long-range correlations, which is normally related to critical state dynamics and gives rise to quick neural network reconfiguration (Battaglia et al., 2020; Linkenkaer-Hansen, Nikouline, Palva, & Ilmoniemi, 2001). In addition, previous work has demonstrated an association between brain signal variability and dynamic functional connectivity across brain regions within six intrinsic connectivity networks in healthy young adults (Fu et al., 2017). Specifically, brain signal variability is negatively correlated with within-network dynamic functional connectivity but positively correlated with between-network dynamic functional connectivity (Fu et al., 2017). Brain signal variability and dynamic functional connectivity are also correlated in time in patients with schizophrenia during rest (Fu et al., 2018, Fu et al., 2021). However, the relationship between brain signal complexity and dynamic functional brain networks in healthy adults is still poorly understood.

In the current study, we used a movie watching dataset collected from adults aged 18 to 55 (Naturalistic Neuroimaging Database; Aliko, Huang, Gheorghiu, Meliss, & Skipper, 2020) to examine how signal variability and complexity link to each other, and to elucidate the relationship between brain signal variability and complexity with static and dynamic functional brain network features. We chose movie watching over resting state because recent work suggests that naturalistic stimuli like movies elicit more distinct brain states as well as higher test-retest reliability of state dynamics (Aliko et al., 2020; Kim, Kay, Shulman, & Corbetta, 2018; van der Meer, Breakspear, Chang, Sonkusare, & Cocchi, 2020, Zhang et al., 2022). In addition, this dataset is unique in that functional acquisition time was sufficiently long to estimate MSE at several time scales. We were specifically interested in how regional BOLD signal dynamics relate to measures of integration (centrality) and segregation (clustering) of the region. We constructed a whole brain functional network by computing functional connectivity between all pairs of regions. We used two centrality measures (betweenness centrality (BC) and eigenvector centrality (EC)), and one non-centrality measure (clustering coefficient (CC)) for each brain region to represent static functional brain networks. For dynamic features of functional brain networks, we applied a sliding window approach to the original time series and calculated these graph theory metrics within each window. We measured dynamic features of functional brain networks by computing the Shannon entropy of these metrics across all windows (Ghaderi, Baltaretu, Andevari, Bharmauria, & Balci, 2020). To estimate the relationship between BOLD signal variability, complexity, features of static and dynamic functional brain networks, we first used linear fixed models and third-degree polynomial fits to evaluate the correlation between BOLD SD or MSE with static/dynamic brain network measures. We then used linear and nonlinear models to predict BOLD SD and MSE using BC, EC and CC. Finally, we operationalized the strength of the relationship between SD and MSE with static/dynamic functional brain network features as the accuracy of prediction. Based on previous empirical work using graph theoretical measures (Easson & McIntosh, 2019), we expected to find a positive correlation between BOLD SD with both static and dynamic functional brain network features. We also expected the relationship between BOLD MSE and functional brain network features to be time scale dependent (McDonough & Nashiro, 2014). Lastly, we hypothesized that for both SD and MSE, the relationship would be stronger with dynamic as compared to static functional brain network measures, because static measures do not reflect the dynamic features of the brain.

## 2 Methods

### 2-1 Participants and study procedure

We used fMRI data from 44 participants (age range: 18-55 yrs, mean age: 25.66 ± 8.37 yrs, 23 women) from the Naturalistic Neuroimaging Database (Aliko et al., 2020), publicly available at https://openneuro.org/datasets/ds002837/versions/2.0.0. All participants were native English speakers, without hearing impairments, and with normal or corrected to normal vision. Other inclusion criteria were being right-handed, having no history of claustrophobia or psychiatric/neurological illness, and currently not taking medication. All participants provided written informed consent and received monetary compensation at the end of the study. The study was approved by the ethics committee of University College London.

### 2-2 MRI data acquisition and preprocessing

Each participant chose one of 10 full-length movies to watch during fMRI scanning, and watched the whole movie in the scanner. Figure 1 shows the number of participants who watched each of the 10 movies. Functional MRI (fMRI) was acquired via a 1.5 T Siemens MAGNETOM Avanto with 32 channel head coil (Siemens Healthcare, Erlangen, Germany), using multiband EPI (TR = 1000 msec; TE = 54.8 msec; flip angle = 75° slices = 40; resolution = 3.2 mm^3^; multiband factor = 4). T1-weighted MPRAGE anatomicals were acquired after fMRI scans (TR = 2730 msec; TE = 3.57 msec; slices = 176; resolution = 1.0 mm^3^). Although previous variability and complexity work using fMRI has been conducted at 3 T, we opted to use this dataset from a 1.5 T scanner because the functional scans were exceptionally long, allowing us to calculate multiple MSE scales. Theoretically, signal to noise ratio (SNR) would double if the data were collected from 3 T. However, a systematic review on direct comparisons between 1.5 and 3 T MRI showed that increases in SNR were only around 25%, and 3 T had worse susceptibility artefacts (Wardlaw et al., 2012).

We used preprocessed fMRI data provided by Aliko et al. (2020; available at https://openneuro.org/datasets/ds002837/versions/2.0.0). Preprocessing steps included slice timing correction, despiking, realignment, coregistration, and spatial normalization to MNI space using AFNI software (Cox, 1996). The time series were further cleaned by detrending and independent component analysis (ICA). We used the Anatomical Automatic Labeling 2 brain atlas (AAL2; Rolls, Joliot, & Tzourio-Mazoyer, 2015) to define 94 regions of interest (ROIs) excluding the cerebellum. We chose unsmoothed data for ROI-based functional connectivity analyses because previous research suggests that smoothing affects node centrality measures of the brain functional network by increasing eigenvector centrality and altering the “hubness” of some nodes (Alakörkkö, Saarimäki, Glerean, Saramäki, & Korhonen, 2017). Specifically, we selected 1800TR (30 min) of continuous data that was free of outliers and excessive motion from each participant. For each participant, we extracted mean time series from each ROI for the estimation of BOLD signal variability, complexity, and FC and dFC measures.

### 2-3 Estimation of BOLD signal variability

We used the standard deviation (SD) of BOLD signal to measure BOLD variability and computed these values in line with previous studies examining BOLD variability (Garrett et al., 2010, 2011; Protzner et al., 2013; Wang, et al., 2021). For each participant, we first normalized 94 ROI time series so that the overall four-dimensional mean was 100. We then divided the data into 90 blocks consisting of 20 TRs (20 sec per block). This block length was previously used in a BOLD SD study for moving-watching data (Wang et al., 2021 supplementary material), which showed that BOLD SD calculation was not affected by block length and block number. We used this same block length for our sliding window size for the estimation of dynamic FC because previous studies suggest that temporal variations in FC are stable when the sliding time window size is between 20 and 40 seconds (Fu et al., 2017; Li et al., 2014). Next, for each ROI, we subtracted the block mean and concatenated data across blocks. We finally calculated ROI-wise SD across this concatenated mean-block corrected time series.

### 2-4 Estimation of BOLD signal complexity

We used MSE to quantify BOLD signal complexity. We calculated MSE using the algorithm available at https://archive.physionet.org/physiotools/mse/tutorial/ (Costa et al., 2002; 2005; Goldberger et al., 2000). The MSE method calculates sample entropy as a measure of predictability (regularity) of brain signal at different timescales, where greater MSE values reflect less predictability or more information content in the signal. To calculate MSE, for each participant and each ROI, the original time series with *N* data points was first downsampled to construct multiple time scales. For a given time scale *τ*, data points within non-overlapping windows of length *τ* of the original time series were averaged to create a data point 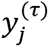 according to equation (1). Thus, 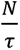 the length of the downsampled time series in scale *τ*. Note that *y*_(1)_ is the original time series.

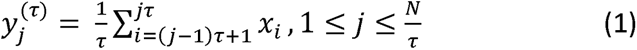

Second, for each time scale, we calculated sample entropy of the corresponding time series by evaluating the appearance of repetitive patterns according to equation (2),

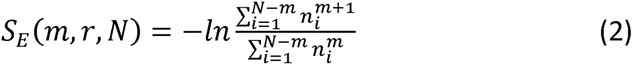

where *N* is the length of the time series in the corresponding time scale, *m* is the pattern length (Small & Tse, 2004), *r* is the similarity criterion (Richman & Moorman, 2000). Sample entropy *S_E_* is the negative natural logarithm of a conditional probability, which estimates the likelihood of two sequences being similar for *m+1* consecutive data points when they are similar for *m* consecutive data points (Costa et al., 2005). Following parameter values used in previous fMRI studies, we set *m* = 2 and *r* = 0.5 for MSE analyses (McDonough & Nashiro, 2014; Smith, Yan, & Wang, 2014; Sokunbi et al., 2013; Wang et al., 2018). The length of each ROI time series was 1800 data points. To ensure reliable MSE estimation, we included only those timescales for which we had at least 50 samples. Thus, for each ROI time series, MSE estimates were sample entropy measures for scales 1 - 36 (or 1 - 36 sec windows), where lower scale values represented fine timescales, and higher scale values represented coarse timescales.

### 2-5 Static and dynamic functional brain networks and nodal network measures

As stated in section 2.3, for each participant, we first divided each of 94 ROI time series into 90 blocks consisting of 20 TRs (20 sec per block). To construct dynamical functional brain networks, we considered each ROI as a network node and the correlation coefficients between mean time series (consisting of 20 TRs) of each pair of ROI regions as network edge. These sets of nodes and edges were then arranged in 94 × 94 adjacency matrices. In each of the 90 adjacency matrices, ROIs were represented in rows and columns, and then functional connectivity between ROIs was represented in corresponding arrays in the adjacency matrices (for each participant and each block, we constructed one adjacency matrix).

We measured three nodal network features: betweenness centrality (BC), eigenvector centrality (EC), and clustering coefficient (CC). *Betweenness centrality (BC)* for a given node is the number of shortest paths between all other nodes that pass the given node (Rubinov & Sporns, 2010). Nodes with higher *BC* provide pathways for fast communication in the networks and can be considered as brain hubs (Sporns, Honey, & Kötter, 2007). Removing a node with high *BC* increases the average length of shortest path (i.e., cost of wiring) in the network (Boccaletti, Latora, Moreno, Chavez, & Hwang, 2006), and then decreases the ability of network to integrate neural signals (Ghaderi, Andevari, & Sowman, 2018)*. Eigenvector centrality (EC)* is defined based on eigenvalue equation of the adjacency matrix (Bonacich, 2007):

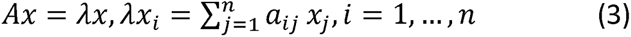

where *A*is adjacency matrix, *λ* is the largest eigenvalue of *A*, *x* is eigenvector associated with *λ*, and *n* is the network size (number of nodes). In equation (3), arrays in the eigenvector *x* represent eigenvector centrality for the nodes. In a weighted undirected network, eigenvector centrality for a given node is associated with connectivity strength for that node with its neighbors and the connectivity strength of those neighbors with their neighbors. Nodes that are connected to other high degree nodes exhibit high eigenvector centrality (Ruhnau, 2000). Akin to *betweenness centrality*, nodes with high values of *EC* can be considered as hubs in the networks (GeethaRamani & Sivaselvi, 2014; Joyce, Laurienti, Burdette, & Hayasaka, 2010). *Clustering coefficient (CC)* is associated with connectivity strength between a given node and its neighbors. A node with high *CC* is strongly connected to its neighbors and those neighbors are strongly connected to each other. The value of *CC* is reduced when connectivity between the node and its neighbors and/or connectivity between the neighbors becomes weak. In a weighted undirected network, *CC* of a given node equals:

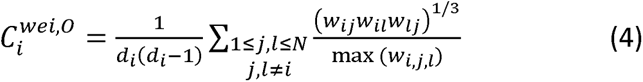

where *d_i_* stands for the summation of connectivity weights of a given node *‘i’*, *w_ij_* is the connectivity weight between the given node ‘*i’* and a neighbor node *‘j’*, and max(*W_i,j,l_*) is the maximum weight between neighbor nodes that make a triangle (Saramäki et al., 2007).

We used the brain connectivity toolbox (Rubinov & Sporns, 2010) in MATLAB (2019b) (at https://www.mathworks.com/products/matlab.html) to calculate these three network measures. To evaluate dFC, we calculated the Shannon entropy for each of these three nodal measures across 90 time points. Mathematically Shannon entropy is defined by (Shannon & Weaver, 1949):

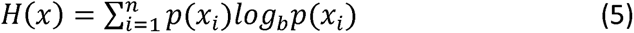

Where *x_i_* are different states, *p* is the occurrence probability of *x_i_* and *n* is the number of states. Minimum Shannon entropy is achieved when the measured values of a given parameter (here *BC, EC, CC*) stay fixed across time (here 90 time-steps). Shannon entropy increases when the distribution of values is random, but a complex or chaotic order of values may also generate higher Shannon entropy (Ghaderi et al., 2020).

### 2-6 Data analysis

#### 2-6-1 Regression analysis

We used linear fixed models to evaluate the correlations between BOLD SD and each scale of MSE. We further estimate the correlations between BOLD SD and MSE with static and dynamic of CC, BC, and EC. For each correlation analysis we reported root mean square error (RMSE), R-squared, t-statistic, and p-value to evaluate the performance of models. The Bonferroni method was used to correct p-values for multiple comparisons. To assess nonlinear relationships between parameters, we also applied third-degree polynomial fits and we calculated RMSE and R-squared for each analysis (because t-statistics and p-values are achieved by linear methods, these values cannot be calculated for the non-linear analyses). We performed these analyses in MATLAB R2019a (https://www.mathworks.com/products/matlab.html).

#### 2-6-2 Linear and nonlinear models

We used six different models including linear (linear and stepwise linear) and nonlinear (tree, support vector machine (SVM), ensemble, and Gaussian) to predict BOLD SD and MSE at each time scale using nodal properties of ROIs. For each model, inputs were the three nodal network measures (i.e., CC, BC, EC) and the output was regional BOLD SD or MSE. To test model accuracy, we used 10-fold cross validation. We further calculated prediction accuracy using Euclidean distance between predicted values and real values:

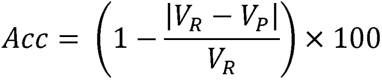

Where *V_R_* is the real value and *V_P_* is the predicted value by model. All these analyses were performed in MATLAB R2019a using the regression learner toolbox (https://www.mathworks.com/products/matlab.html).

## 3 Results

### 3-1 Regression analyses – SD and MSE

Our linear fixed model analysis identified a significant correlation between BOLD SD and MSE at all scales (absolute values ranging from *R^2^* = 0.0906, RMSE = 0.1106, and *p* = 0.003 to values ranging from *R^2^* = 0.5725, *RMSE* = 0.0280, and *p* <0.001). The association was positive between SD and MSE at the first two scales and negative at the remaining coarser scales (Table 4.1).

**Table 4. 1.** Linear regression: Correlations between SD and MSE

|  | **BOLD SD** |  |  |  |
| --- | --- | --- | --- | --- |
| MSE time scale | RMSE | R-squared | t-statistic | p-value |
| 1 | 0.0862 | 0.3804 | 7.516 | <1e-3 |
| 2 | 0.0316 | 0.0906 | 3.0275 | 3.2e-3 |
| 3 | 0.028 | 0.5725 | -11.0991 | <1e-3 |
| 4 | 0.0523 | 0.5556 | -10.7245 | <1e-3 |
| 5 | 0.0682 | 0.5425 | -10.4458 | <1e-3 |
| 6 | 0.0796 | 0.5277 | -10.138 | <1e-3 |
| 7 | 0.0876 | 0.521 | -10.0031 | <1e-3 |
| 8 | 0.0933 | 0.5093 | -9.7717 | <1e-3 |
| 9 | 0.0986 | 0.4961 | -9.5168 | <1e-3 |
| 10 | 0.1024 | 0.4912 | -9.4235 | <1e-3 |
| 11 | 0.105 | 0.4951 | -9.4985 | <1e-3 |
| 12 | 0.1076 | 0.4843 | -9.2951 | <1e-3 |
| 13 | 0.1088 | 0.4816 | -9.2451 | <1e-3 |
| 14 | 0.1105 | 0.4784 | -9.1856 | <1e-3 |
| 15 | 0.1106 | 0.4791 | -9.1993 | <1e-3 |
| 16 | 0.1106 | 0.4766 | -9.1524 | <1e-3 |
| 17 | 0.1093 | 0.4849 | -9.3065 | <1e-3 |
| 18 | 0.1104 | 0.4772 | -9.1642 | <1e-3 |
| 19 | 0.1091 | 0.4702 | -9.0361 | <1e-3 |
| 20 | 0.1083 | 0.4717 | -9.0626 | <1e-3 |
| 21 | 0.1066 | 0.4701 | -9.0336 | <1e-3 |
| 22 | 0.1057 | 0.478 | -9.178 | <1e-3 |
| 23 | 0.1052 | 0.4562 | -8.786 | <1e-3 |
| 24 | 0.1042 | 0.458 | -8.8166 | <1e-3 |
| 25 | 0.101 | 0.458 | -8.8167 | <1e-3 |
| 26 | 0.1011 | 0.4595 | -8.8438 | <1e-3 |
| 27 | 0.098 | 0.4612 | -8.8747 | <1e-3 |
| 28 | 0.0967 | 0.453 | -8.7281 | <1e-3 |
| 29 | 0.0946 | 0.4484 | -8.6485 | <1e-3 |
| 30 | 0.0925 | 0.4502 | -8.6788 | <1e-3 |
| 31 | 0.0899 | 0.4533 | -8.7343 | <1e-3 |
| 32 | 0.0878 | 0.4519 | -8.7097 | <1e-3 |
| 33 | 0.0865 | 0.4472 | -8.6277 | <1e-3 |
| 34 | 0.0865 | 0.4501 | -8.6783 | <1e-3 |
| 35 | 0.0831 | 0.4391 | -8.4869 | <1e-3 |
| 36 | 0.0828 | 0.4351 | -8.4178 | <1e-3 |

### 3-2 Regression analyses – SD and MSE with static nodal measures

We used linear fixed models and third-degree polynomial fits to evaluate the relationship between static nodal brain network measures with BOLD SD and MSE, separately. We found significant negative correlations between nodal network measures and SD (see Table 4.2, Figures 4.2 and 4.3). These correlations were higher when we applied third-degree polynomial fits, which suggested that there was a nonlinear relationship between SD and static nodal measures (the best fit was for CC: *R^2^* = 0.657, *RMSE* = 0.0014; see Table 4.2).

**Figure 4. 1.**
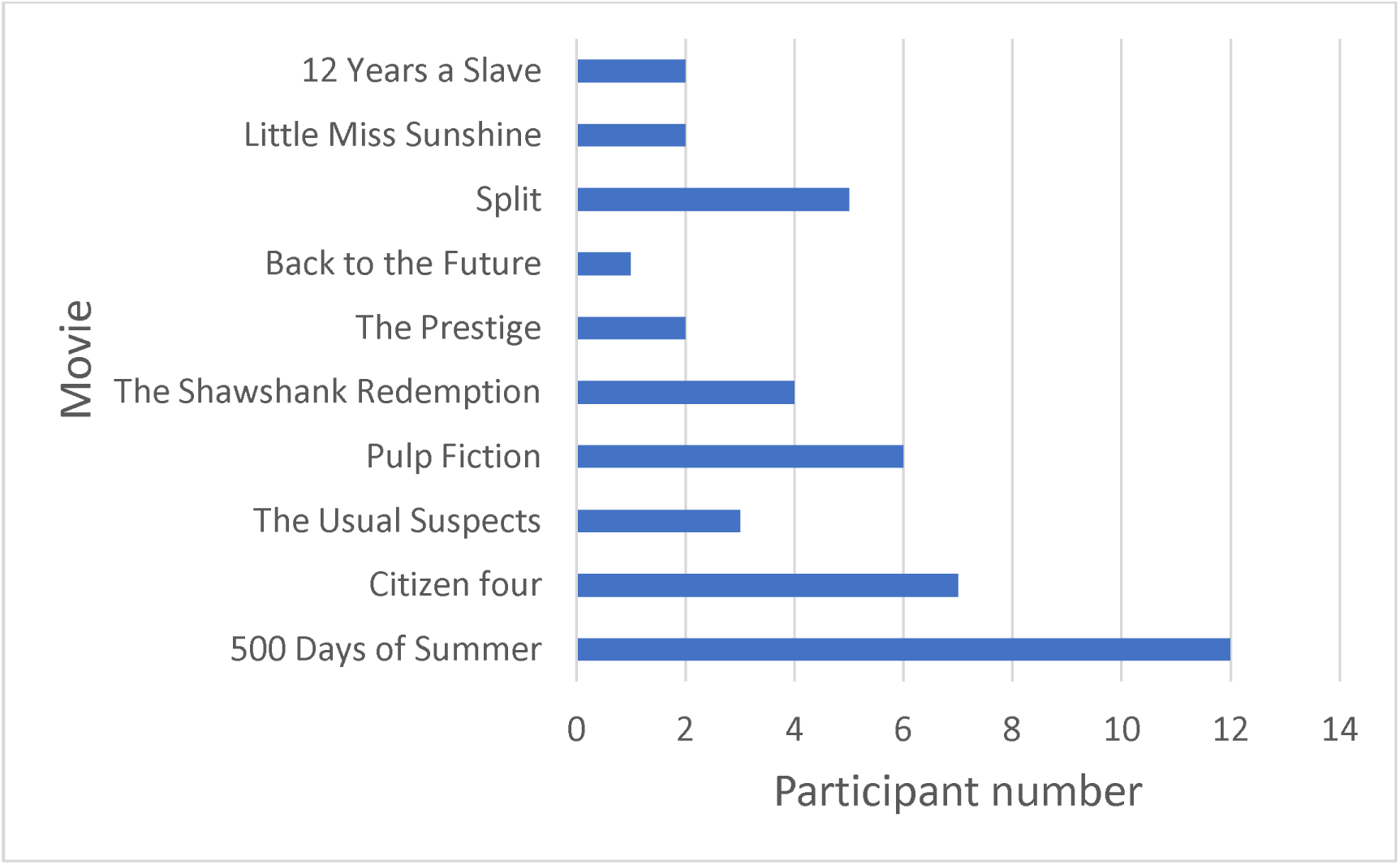
Number of participants that watched each of the 10 movies.

**Figure 4. 2.**
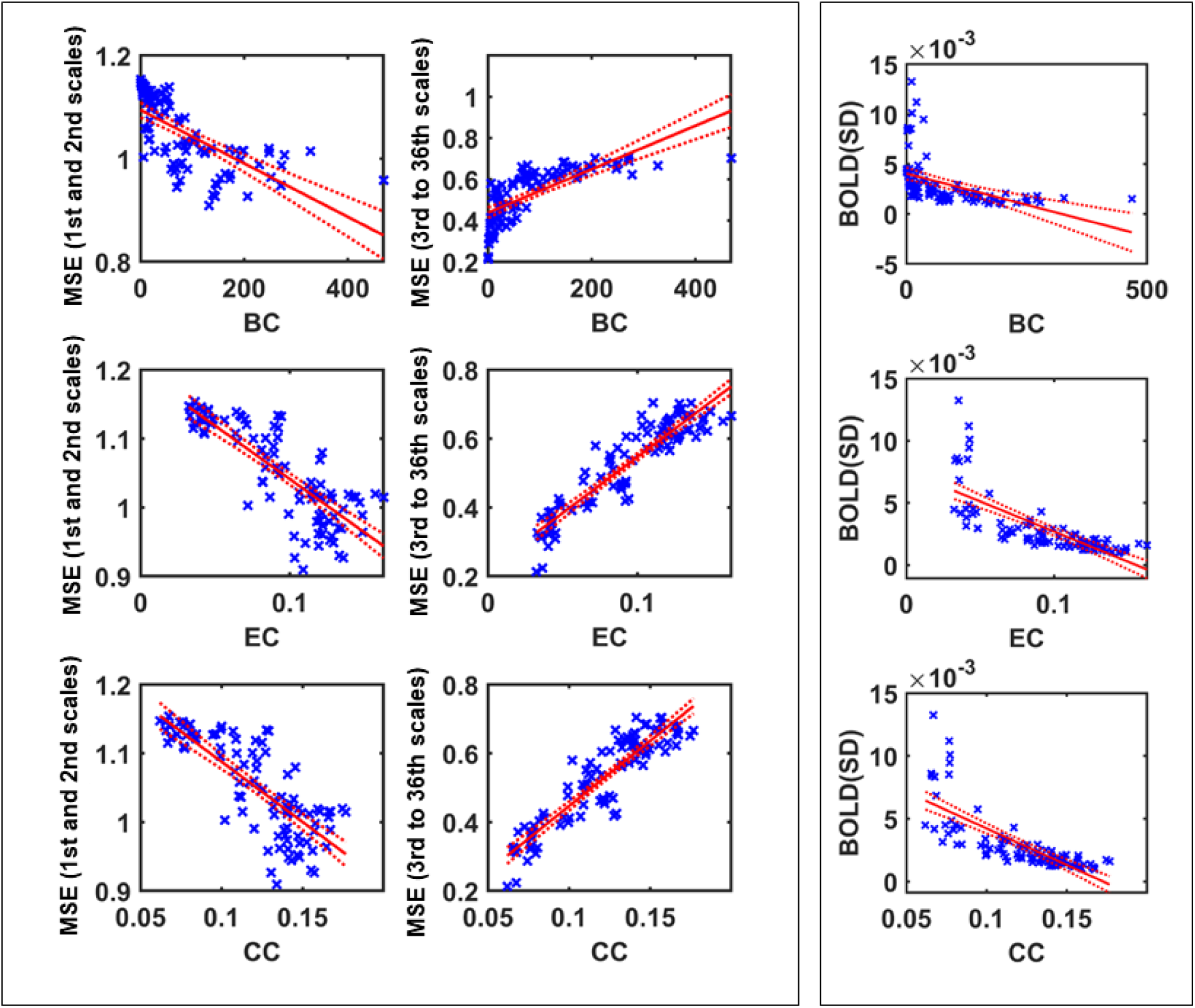
Linear fixed model regression results between MSE (left), BOLD SD (right) and static nodal measures. For visualization of all scales of MSE, we averaged over all negative correlations (1^st^ and 2^nd^ scales) and all positive correlations (3rd to 36th scales) separately, in different plots. Each point of scatter plot stands for one brain region.

**Table 4. 2.**
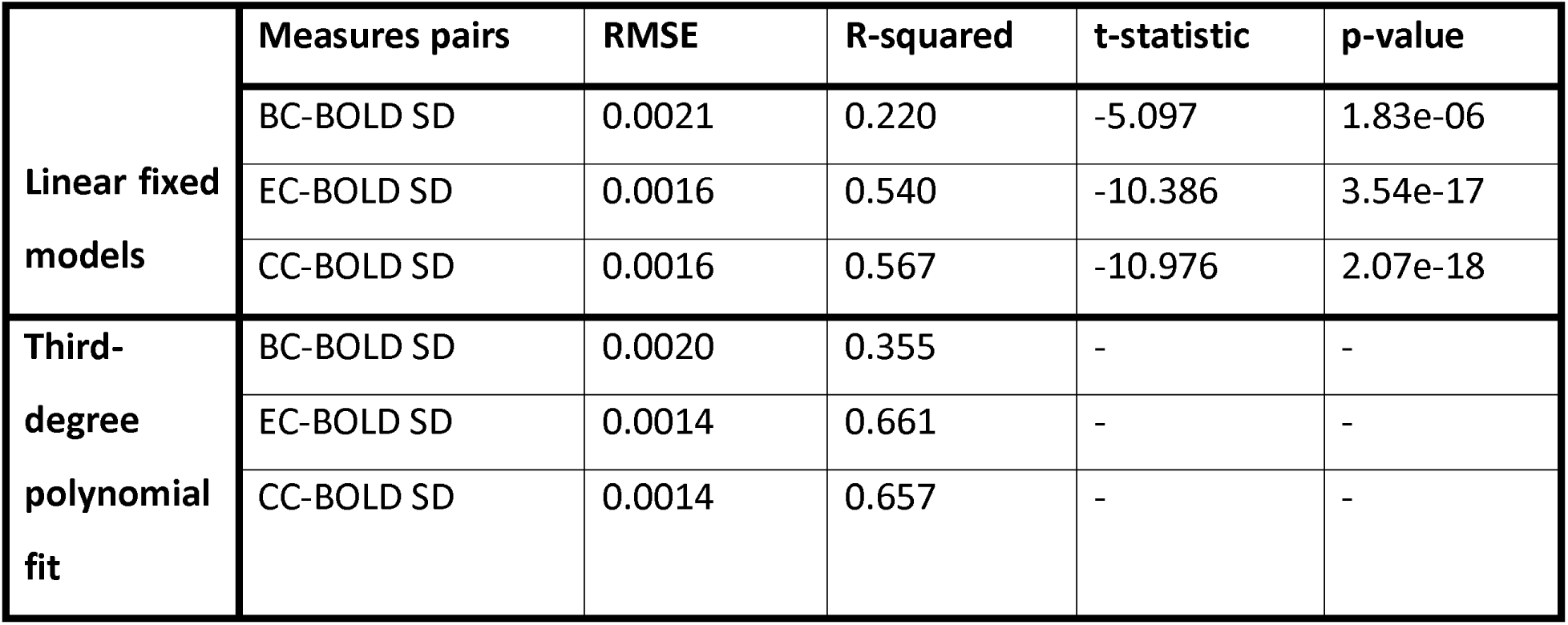
Correlations between SD and static nodal network features

|  | Measures pairs | RMSE | R-squared | t-statistic | p-value |
| --- | --- | --- | --- | --- | --- |
| Linear fixed models | BC-BOLD SD | 0.0021 | 0.220 | -5.097 | 1.83e-06 |
|  | EC-BOLD SD | 0.0016 | 0.540 | -10.386 | 3.54e-17 |
|  | CC-BOLD SD | 0.0016 | 0.567 | -10.976 | 2.07e-18 |
| Third-degree polynomial fit | BC-BOLD SD | 0.0020 | 0.355 | - | - |
|  | EC-BOLD SD | 0.0014 | 0.661 | - | - |
|  | CC-BOLD SD | 0.0014 | 0.657 | - | - |

We also found that static nodal network measures (BC, EC, and CC) were negatively correlated with MSE at the first and second scales but positively correlated with the remaining coarser scales (the absolute values obtained from linear regression models ranged between *R^2^* = 0.1879, *RMSE* = 0.1079, *p*<0.001 and *R^2^* = 0.8469, *RMSE* = 0.0253, *p*<0.001; see Table 4.3, Figures 4.2 and 4.3). The associations between BC and MSE at all scales were improved with third-degree polynomial fits which suggested that the relationship between MSE and BC was nonlinear (see Table 4.4).

**Figure 4. 3.**
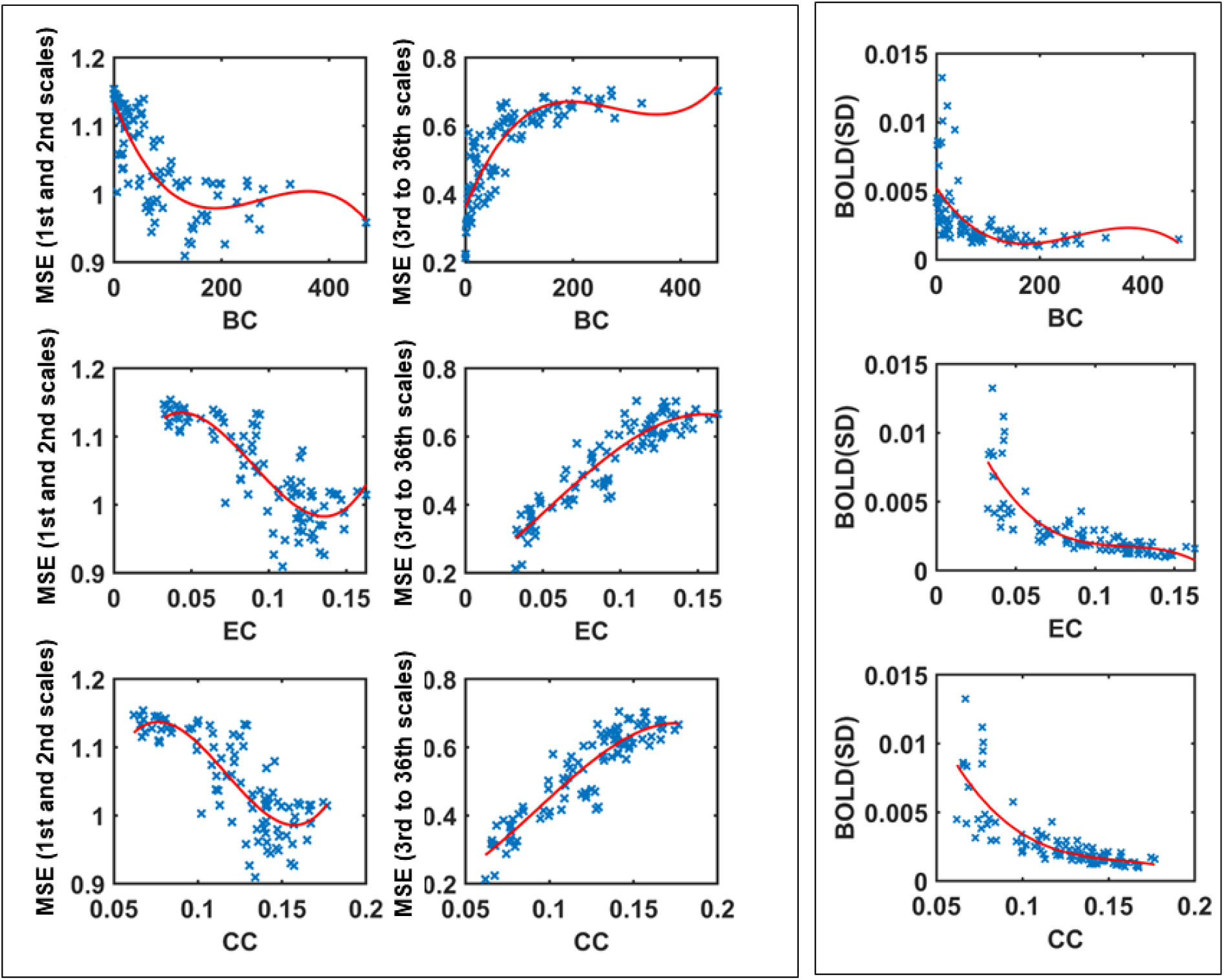
Third degree polynomial regression results between MSE (left), BOLD SD (right) and static nodal measures. For visualization of all scales of MSE, we averaged over all negative correlations (1^st^ and 2^nd^ scales) and all positive correlations (3rd to 36th scales) separately, in different plots. Each point of scatter plot stands for one brain region. Each point of scatter plot stands for one brain region.

**Table 4. 3.**
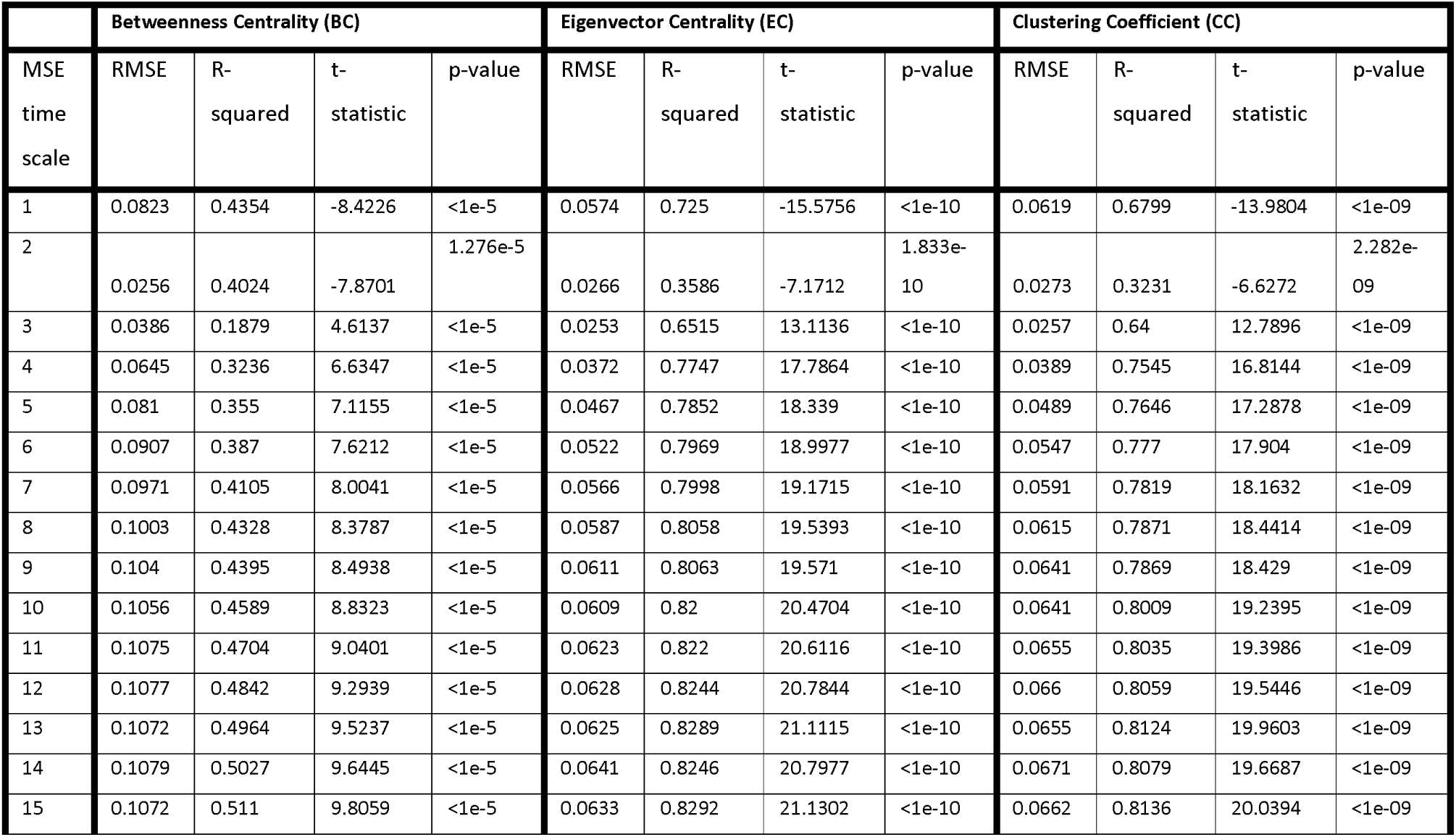
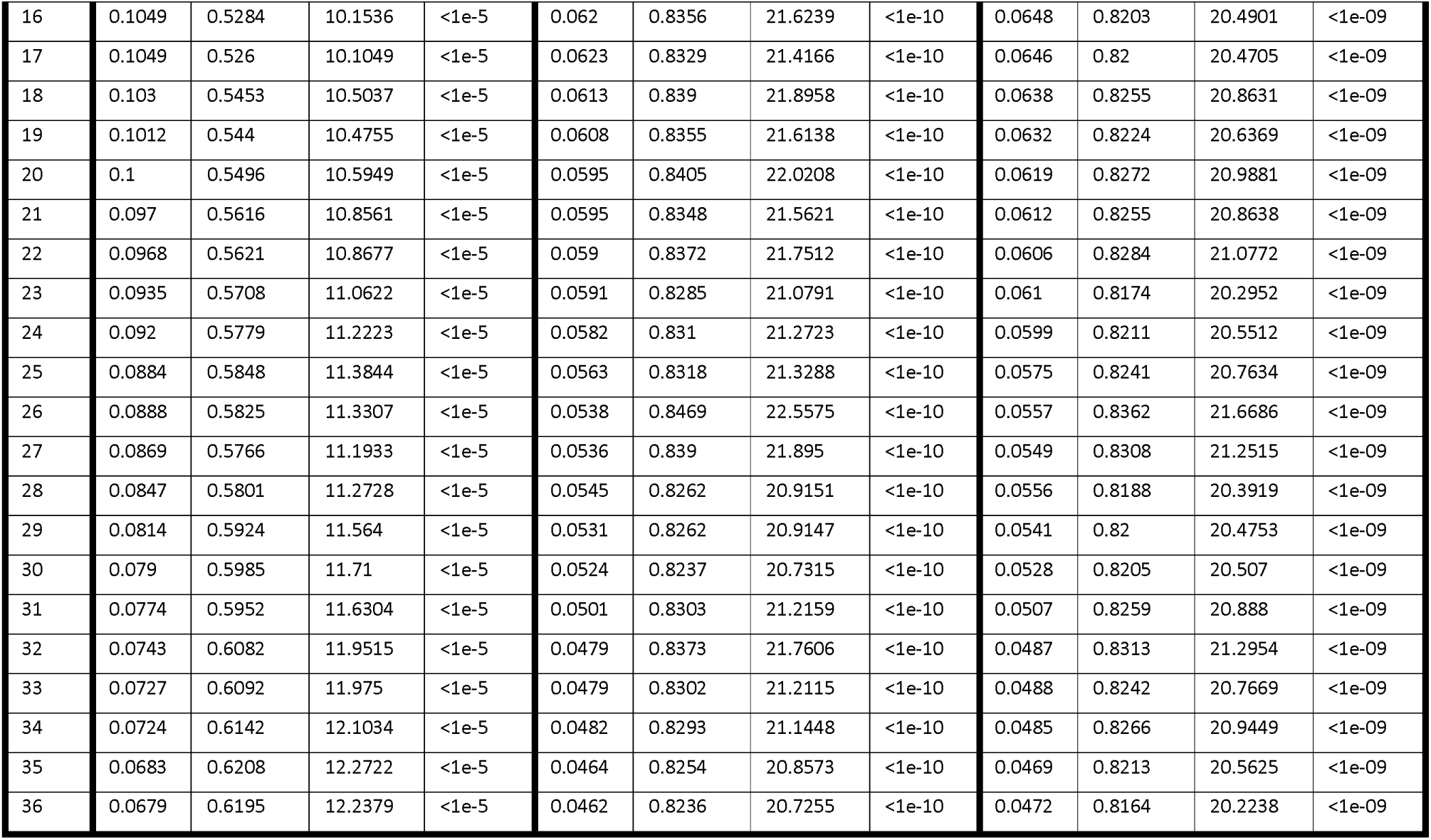
Linear regression. Correlations between MSE and static nodal network features

**Table 4. 4.**
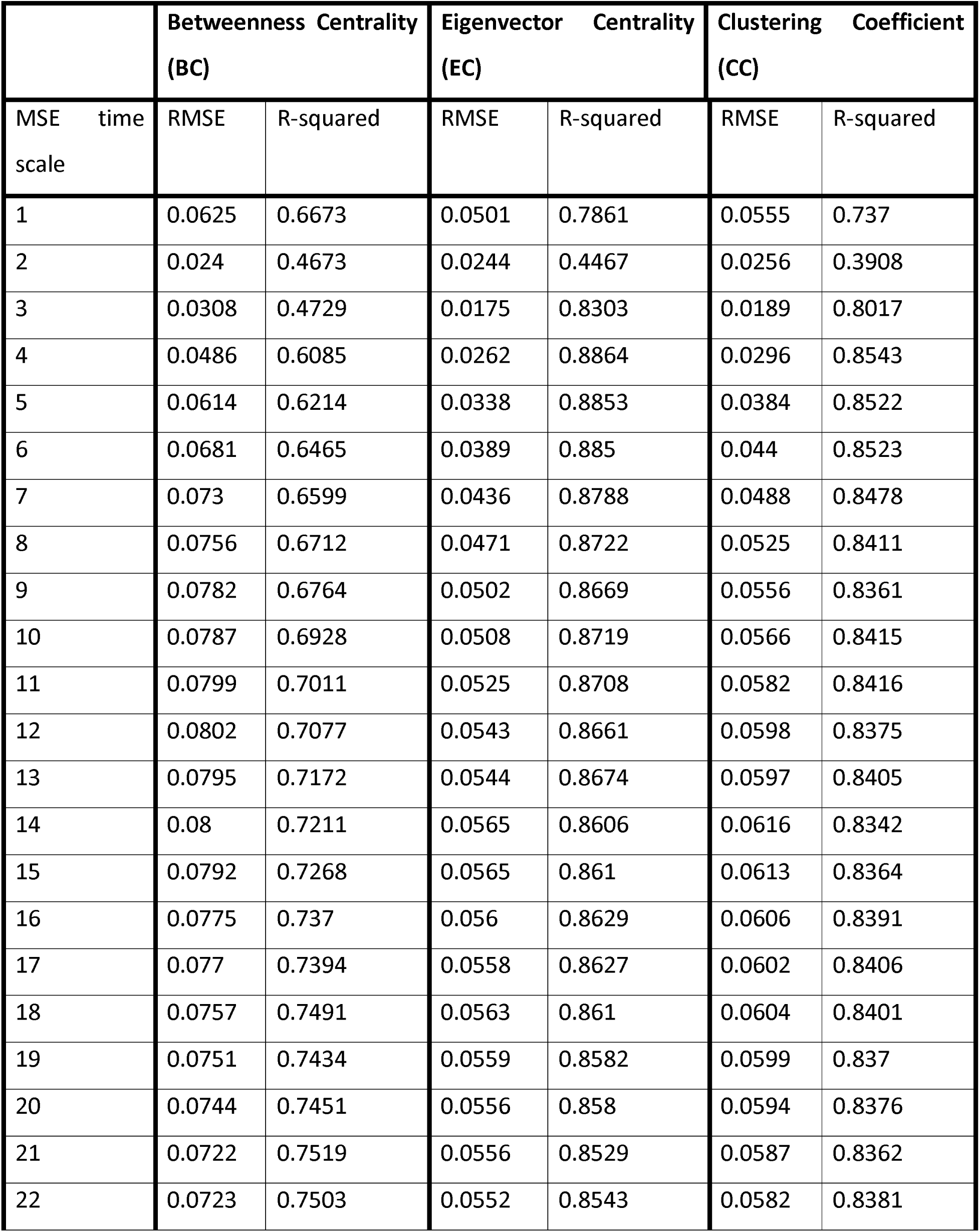
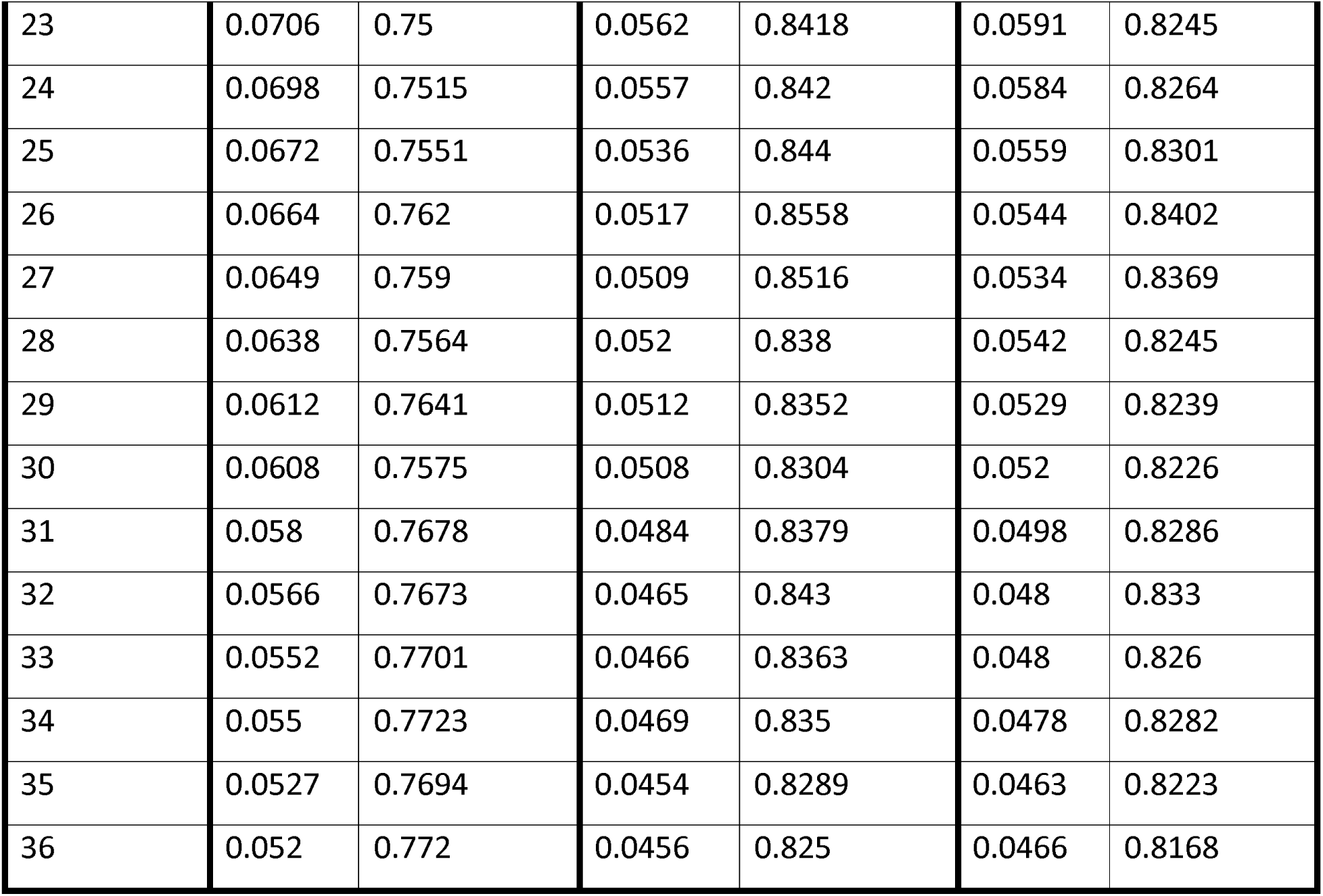
Third-degree polynomial fit. Correlations between MSE and *static* nodal network features

|  | Betweenness Centrality (BC) |  | Eigenvector Centrality (EC) |  | Clustering Coefficient (CC) |  |
| --- | --- | --- | --- | --- | --- | --- |
| MSE time<br>scale | RMSE | R-squared | RMSE | R-squared | RMSE | R-squared |
| 1 | 0.0625 | 0.6673 | 0.0501 | 0.7861 | 0.0555 | 0.737 |
| 2 | 0.024 | 0.4673 | 0.0244 | 0.4467 | 0.0256 | 0.3908 |
| 3 | 0.0308 | 0.4729 | 0.0175 | 0.8303 | 0.0189 | 0.8017 |
| 4 | 0.0486 | 0.6085 | 0.0262 | 0.8864 | 0.0296 | 0.8543 |
| 5 | 0.0614 | 0.6214 | 0.0338 | 0.8853 | 0.0384 | 0.8522 |
| 6 | 0.0681 | 0.6465 | 0.0389 | 0.885 | 0.044 | 0.8523 |
| 7 | 0.073 | 0.6599 | 0.0436 | 0.8788 | 0.0488 | 0.8478 |
| 8 | 0.0756 | 0.6712 | 0.0471 | 0.8722 | 0.0525 | 0.8411 |
| 9 | 0.0782 | 0.6764 | 0.0502 | 0.8669 | 0.0556 | 0.8361 |
| 10 | 0.0787 | 0.6928 | 0.0508 | 0.8719 | 0.0566 | 0.8415 |
| 11 | 0.0799 | 0.7011 | 0.0525 | 0.8708 | 0.0582 | 0.8416 |
| 12 | 0.0802 | 0.7077 | 0.0543 | 0.8661 | 0.0598 | 0.8375 |
| 13 | 0.0795 | 0.7172 | 0.0544 | 0.8674 | 0.0597 | 0.8405 |
| 14 | 0.08 | 0.7211 | 0.0565 | 0.8606 | 0.0616 | 0.8342 |
| 15 | 0.0792 | 0.7268 | 0.0565 | 0.861 | 0.0613 | 0.8364 |
| 16 | 0.0775 | 0.737 | 0.056 | 0.8629 | 0.0606 | 0.8391 |
| 17 | 0.077 | 0.7394 | 0.0558 | 0.8627 | 0.0602 | 0.8406 |
| 18 | 0.0757 | 0.7491 | 0.0563 | 0.861 | 0.0604 | 0.8401 |
| 19 | 0.0751 | 0.7434 | 0.0559 | 0.8582 | 0.0599 | 0.837 |
| 20 | 0.0744 | 0.7451 | 0.0556 | 0.858 | 0.0594 | 0.8376 |
| 21 | 0.0722 | 0.7519 | 0.0556 | 0.8529 | 0.0587 | 0.8362 |
| 22 | 0.0723 | 0.7503 | 0.0552 | 0.8543 | 0.0582 | 0.8381 |
| 23 | 0.0706 | 0.75 | 0.0562 | 0.8418 | 0.0591 | 0.8245 |
| 24 | 0.0698 | 0.7515 | 0.0557 | 0.842 | 0.0584 | 0.8264 |
| 25 | 0.0672 | 0.7551 | 0.0536 | 0.844 | 0.0559 | 0.8301 |
| 26 | 0.0664 | 0.762 | 0.0517 | 0.8558 | 0.0544 | 0.8402 |
| 27 | 0.0649 | 0.759 | 0.0509 | 0.8516 | 0.0534 | 0.8369 |
| 28 | 0.0638 | 0.7564 | 0.052 | 0.838 | 0.0542 | 0.8245 |
| 29 | 0.0612 | 0.7641 | 0.0512 | 0.8352 | 0.0529 | 0.8239 |
| 30 | 0.0608 | 0.7575 | 0.0508 | 0.8304 | 0.052 | 0.8226 |
| 31 | 0.058 | 0.7678 | 0.0484 | 0.8379 | 0.0498 | 0.8286 |
| 32 | 0.0566 | 0.7673 | 0.0465 | 0.843 | 0.048 | 0.833 |
| 33 | 0.0552 | 0.7701 | 0.0466 | 0.8363 | 0.048 | 0.826 |
| 34 | 0.055 | 0.7723 | 0.0469 | 0.835 | 0.0478 | 0.8282 |
| 35 | 0.0527 | 0.7694 | 0.0454 | 0.8289 | 0.0463 | 0.8223 |
| 36 | 0.052 | 0.772 | 0.0456 | 0.825 | 0.0466 | 0.8168 |

### 3-3 Regression analyses – SD and MSE with dynamic nodal network measures

We found the highest Shannon entropy values for EC (average over participants: 5.887), and the lowest entropy values for BC (average over participants: 5.259). However, the variation of Shannon entropy across regions was higher in BC than EC and CC (Figure 4.4). Both the linear fixed model and third-degree polynomial fits showed significant negative correlations between SD and the entropy of BC and CC (Table 4.5 and Figure 4.5, 4.6). However, the correlation between SD and the entropy of EC was not significant (Table 4.5).

**Figure 4. 4.**
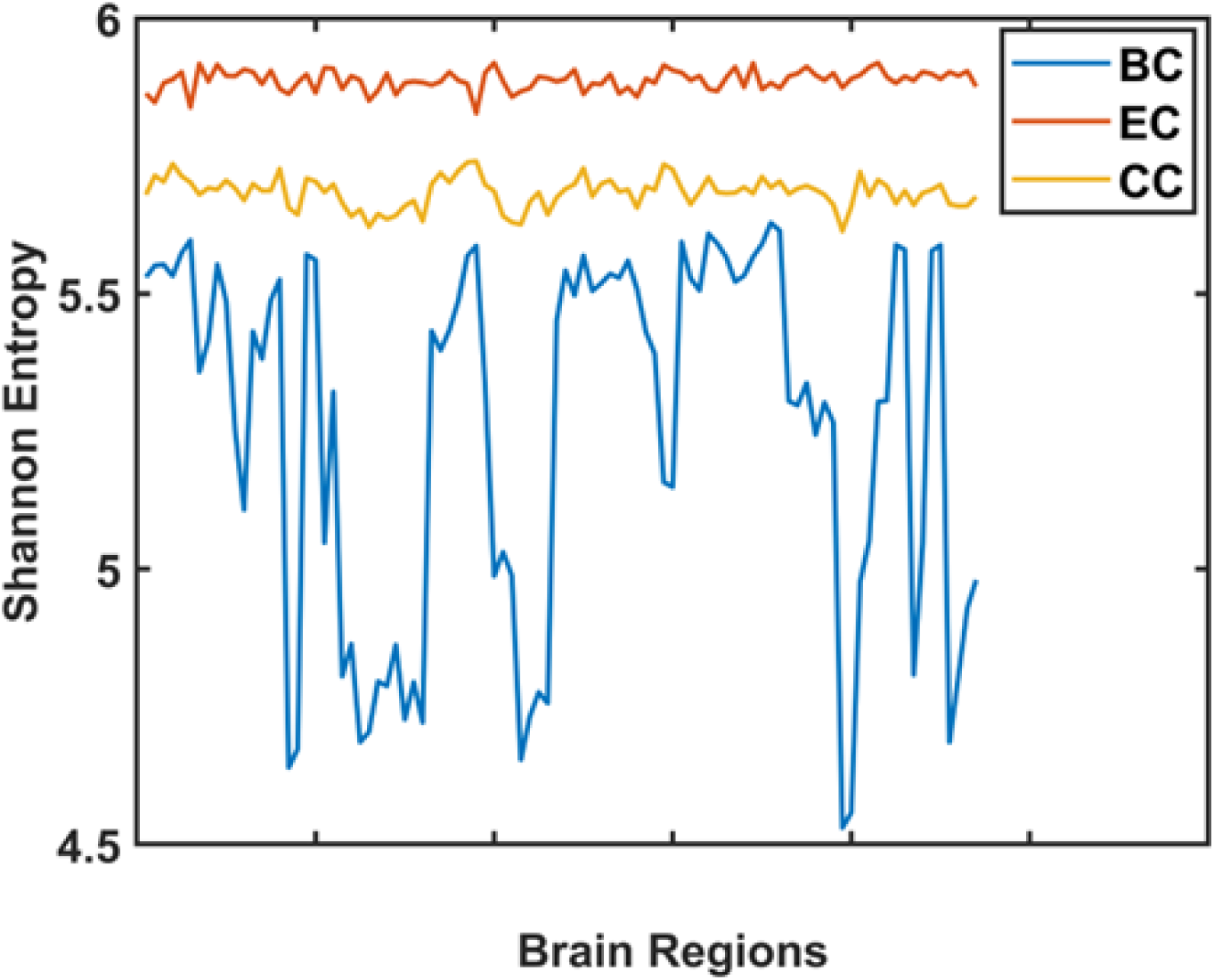
Shannon entropy of nodal measures in different regions (averaged over all participants). EC exhibited highest Shannon entropy, and BC exhibited highest variation of Shannon entropy across regions.

**Table 4. 5.**
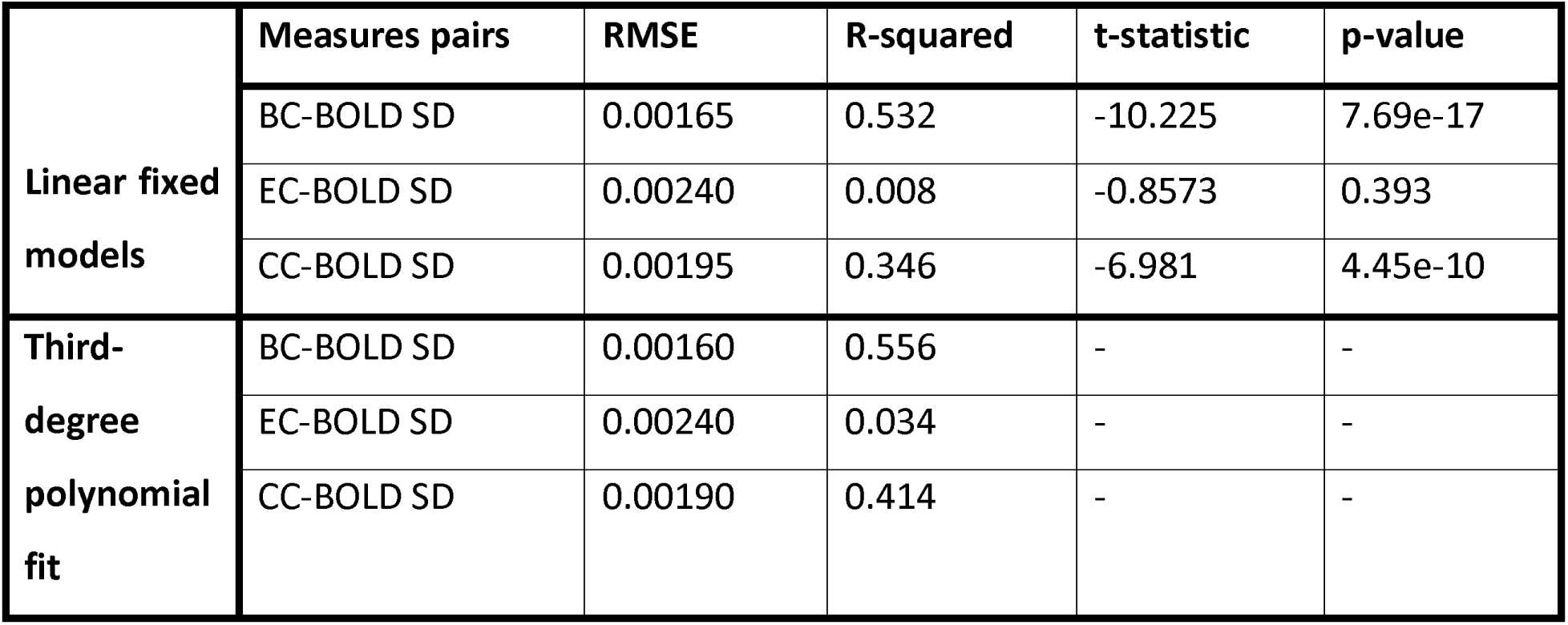
Correlations between BOLD SD and *dynamical* nodal network features (Shannon entropy)

|  | Measures pairs | RMSE | R-squared | t-statistic | p-value |
| --- | --- | --- | --- | --- | --- |
| Linear fixed models | BC-BOLD SD | 0.00165 | 0.532 | -10.225 | 7.69e-17 |
|  | EC-BOLD SD | 0.00240 | 0.008 | -0.8573 | 0.393 |
|  | CC-BOLD SD | 0.00195 | 0.346 | -6.981 | 4.45e-10 |
| Third-degree polynomial fit | BC-BOLD SD | 0.00160 | 0.556 | - | - |
|  | EC-BOLD SD | 0.00240 | 0.034 | - | - |
|  | CC-BOLD SD | 0.00190 | 0.414 | - | - |

Results from our linear and nonlinear regression models between MSE and dynamic nodal measures are shown in Tables 4.6 – 4.7 and Figures 4.5 – 4.6. Both models revealed high correlations between the entropy of BC and MSE (absolute values ranging from *R^2^* = 0.4494, *RMSE* = 0.2497 to *R^2^* = 0.9761, *RMSE* = 0.0515), as well as significant medium to high correlations between the entropy of CC and MSE. These were negative at fine scales (1^st^ and 2^nd^ scales) but positive at coarse scales (3^rd^ to 36^th^ scales; absolute values ranged from *R^2^* = 0.0747, *RMSE* = 0.0528 and *R^2^* = 0.8313, *RMSE* = 0.0199). Finally, correlations between the entropy of EC and MSE for all scales were not significant (see Table 4.6).

**Figure 4. 5.**
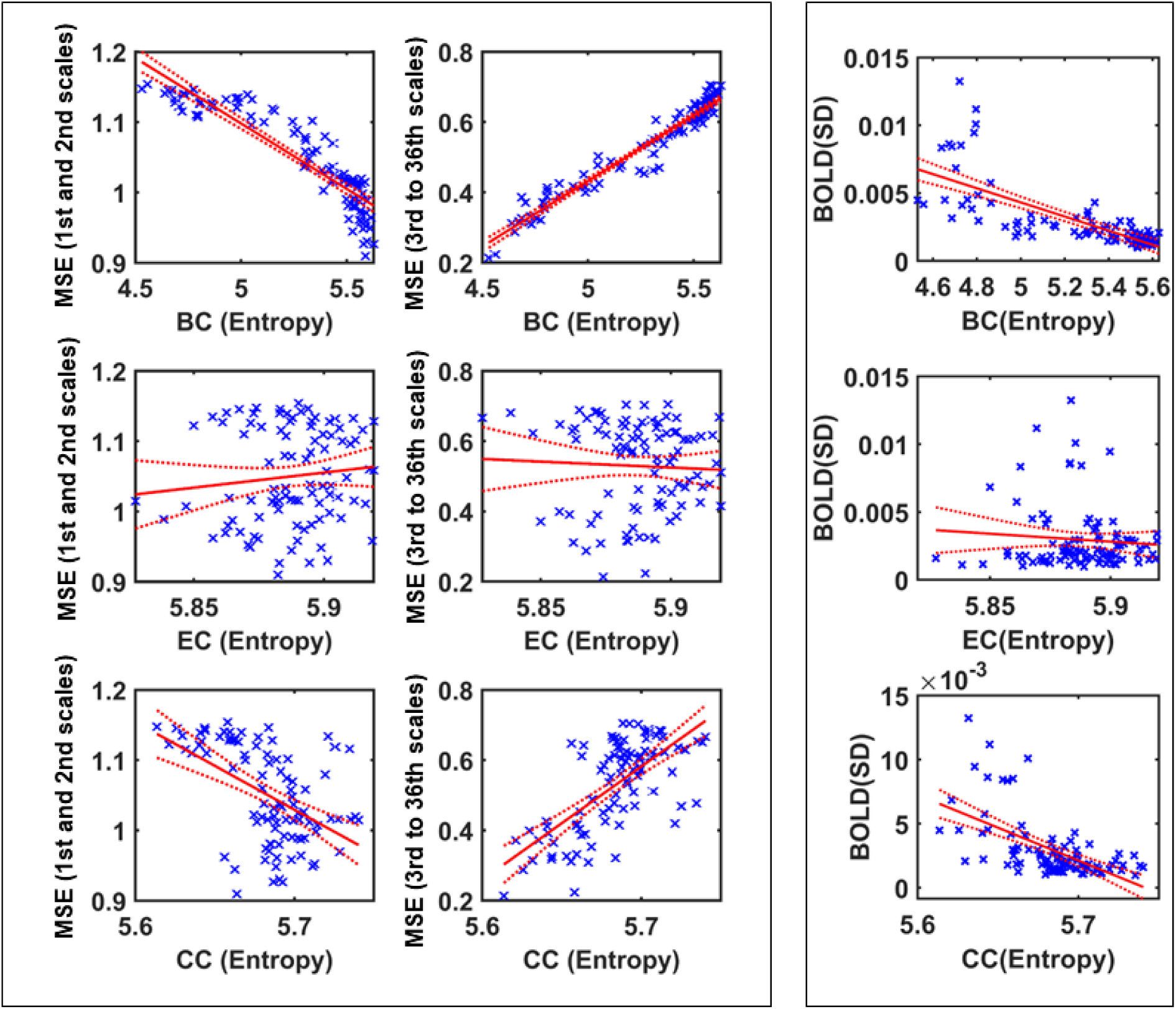
Linear fixed model regression results between MSE (left), BOLD SD (right) and dynamical nodal measures (Shannon entropy of measures during time). For visualization of all scales of MSE, we averaged over all negative correlations (1^st^ and 2^nd^ scales) and all positive correlations (3rd to 36th scales) separately, in different plots. Each point of scatter plot stands for one brain region.

**Figure 4. 6.**
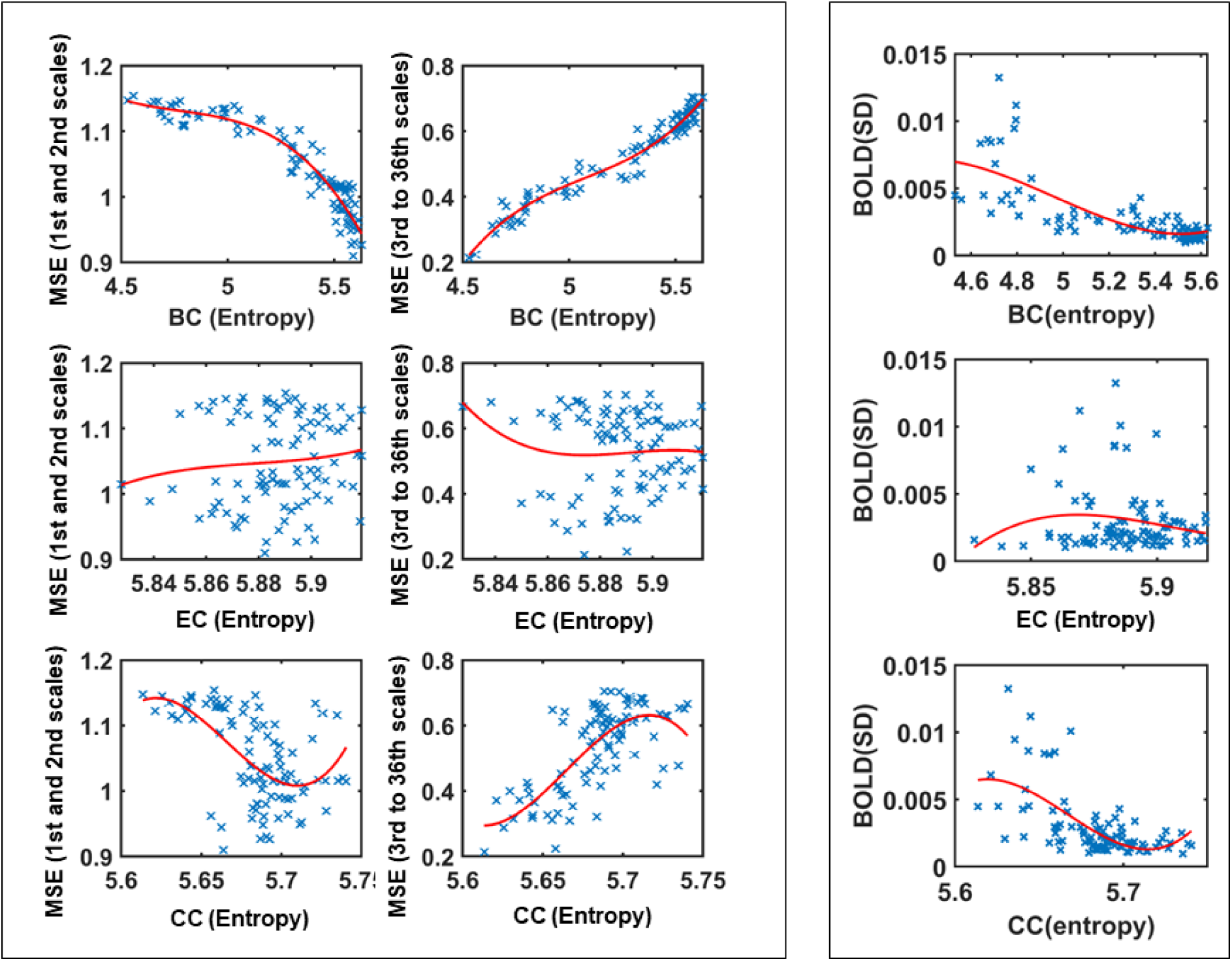
Third degree polynomial regression results between MSE (left), BOLD SD (right) and dynamical nodal measures (entropy of measures during time). For visualization of all scales of MSE, we averaged over all negative correlations (1^st^ and 2^nd^ scales) and all positive correlations (3rd to 36th scales) separately, in different plots. Each point of scatter plot stands for one brain region.

**Table 4. 6.**
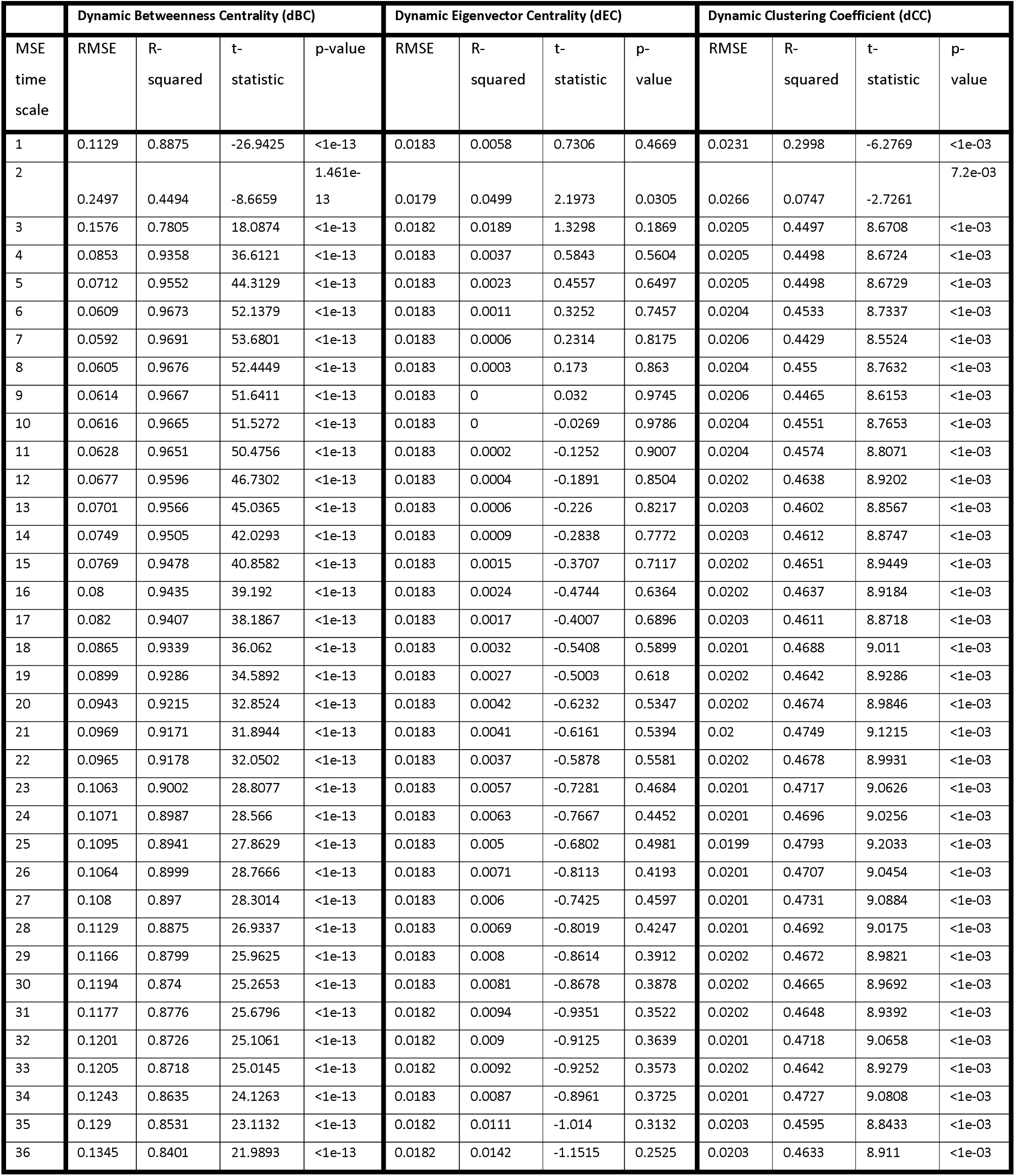
Linear regression. Correlations between MSE and dynamic nodal network features

### 3-4 Prediction of SD and MSE using static nodal network measures

We used six models to predict SD based on static nodal measures (i.e., BC, EC, CC). The accuracy of SD prediction across all models and all regions was higher than 0.78, which is reliable (see Table 4.8, and Figure 4.7). The tree model showed the highest average accuracy across regions of 0.8 (Table 4.8).

**Figure 4. 7.**
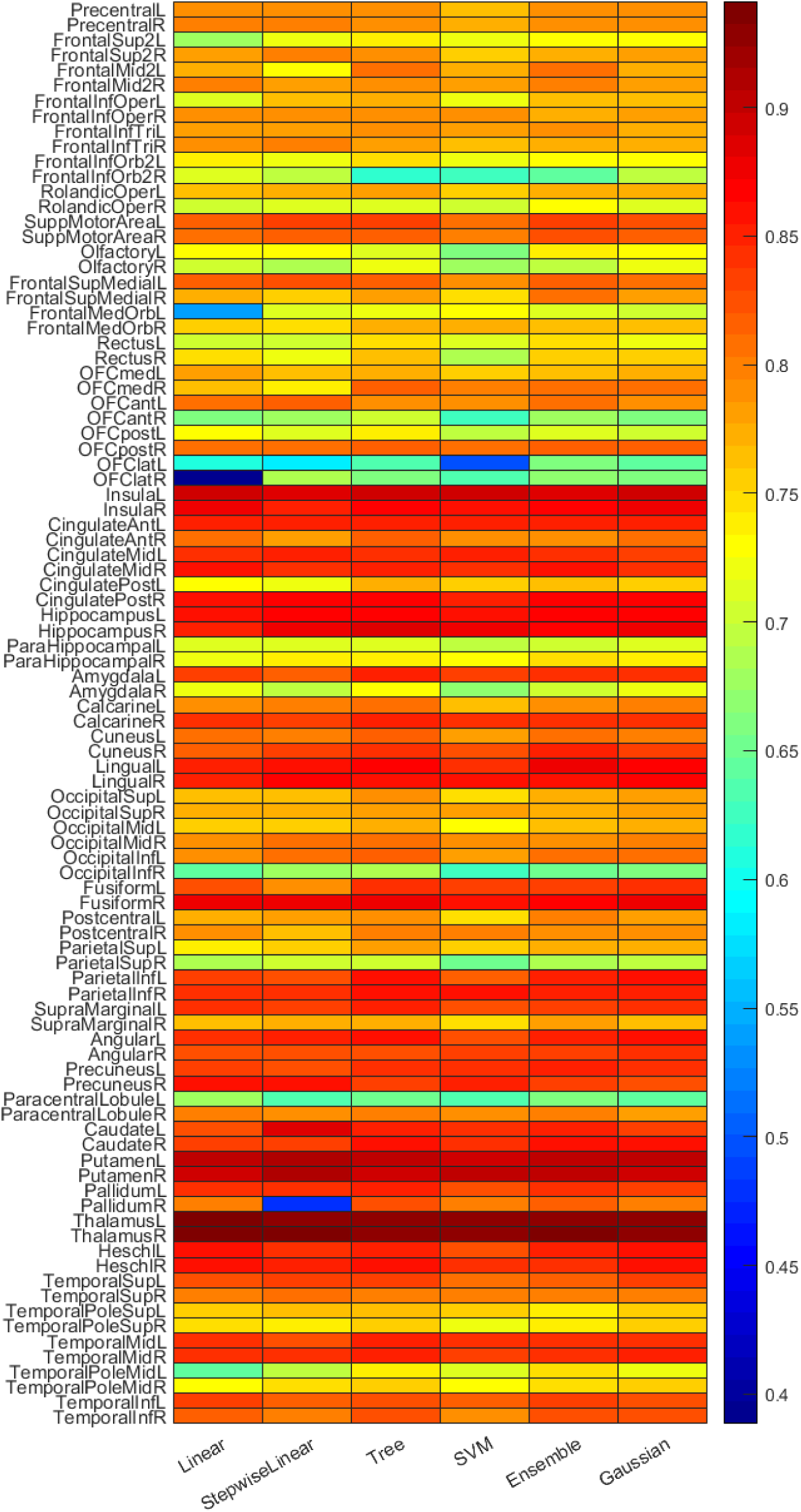
BOLD SD prediction accuracy based on *static* nodal network features (BC, EC, CC).

**Figure 4. 8.**
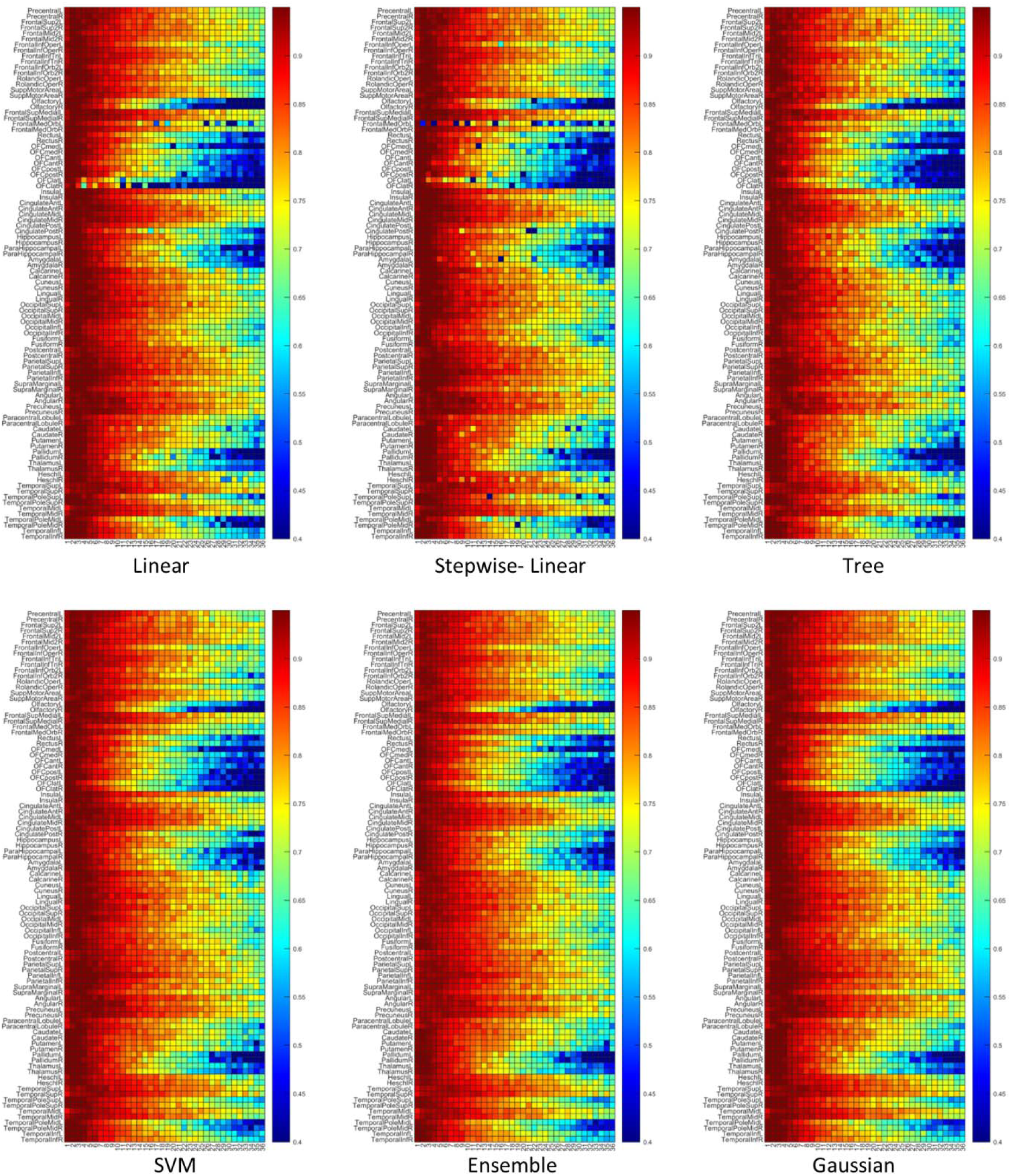
MSE prediction accuracy based on *static* nodal network features (BC, EC, CC).

**Table 4. 7.**
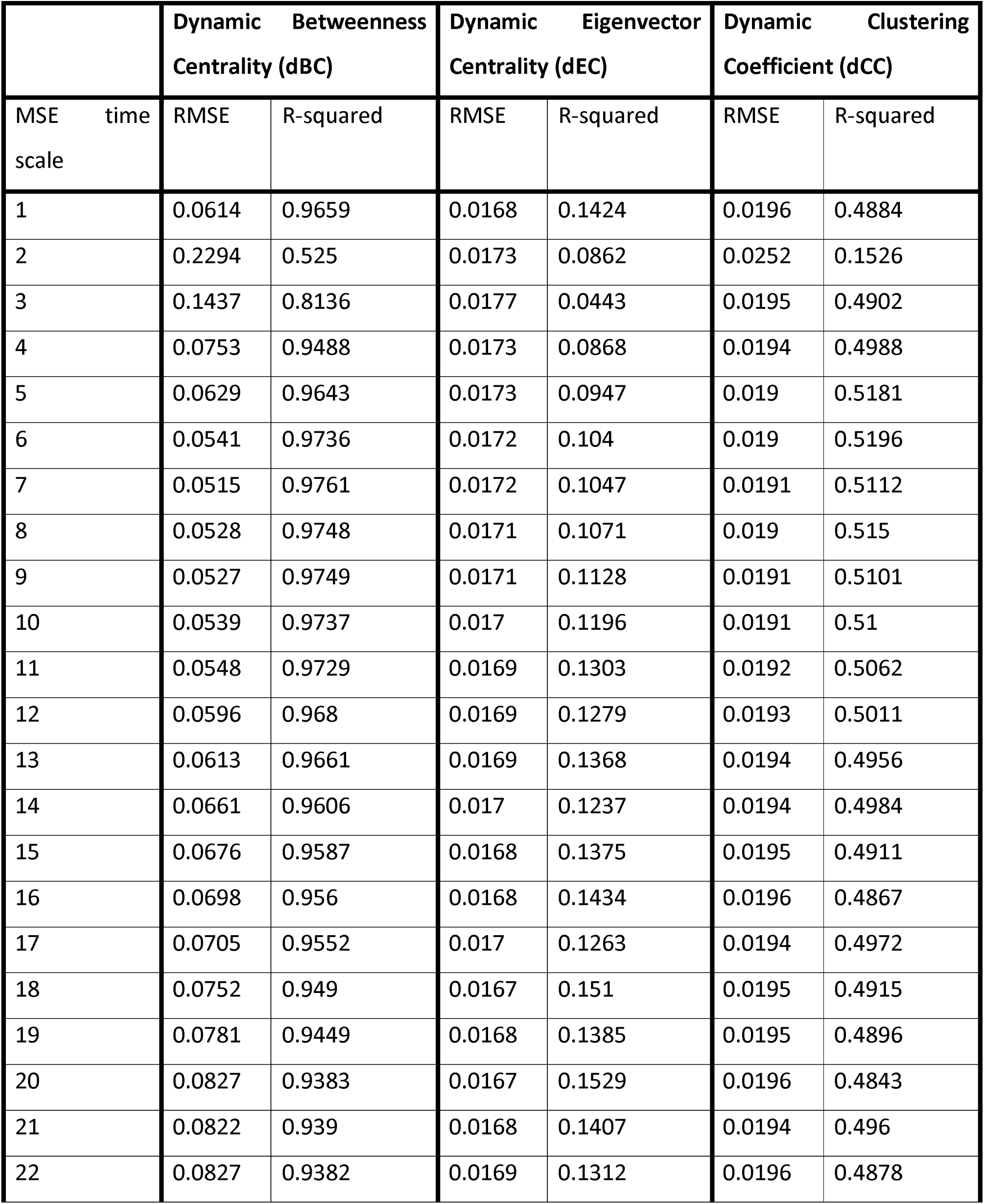
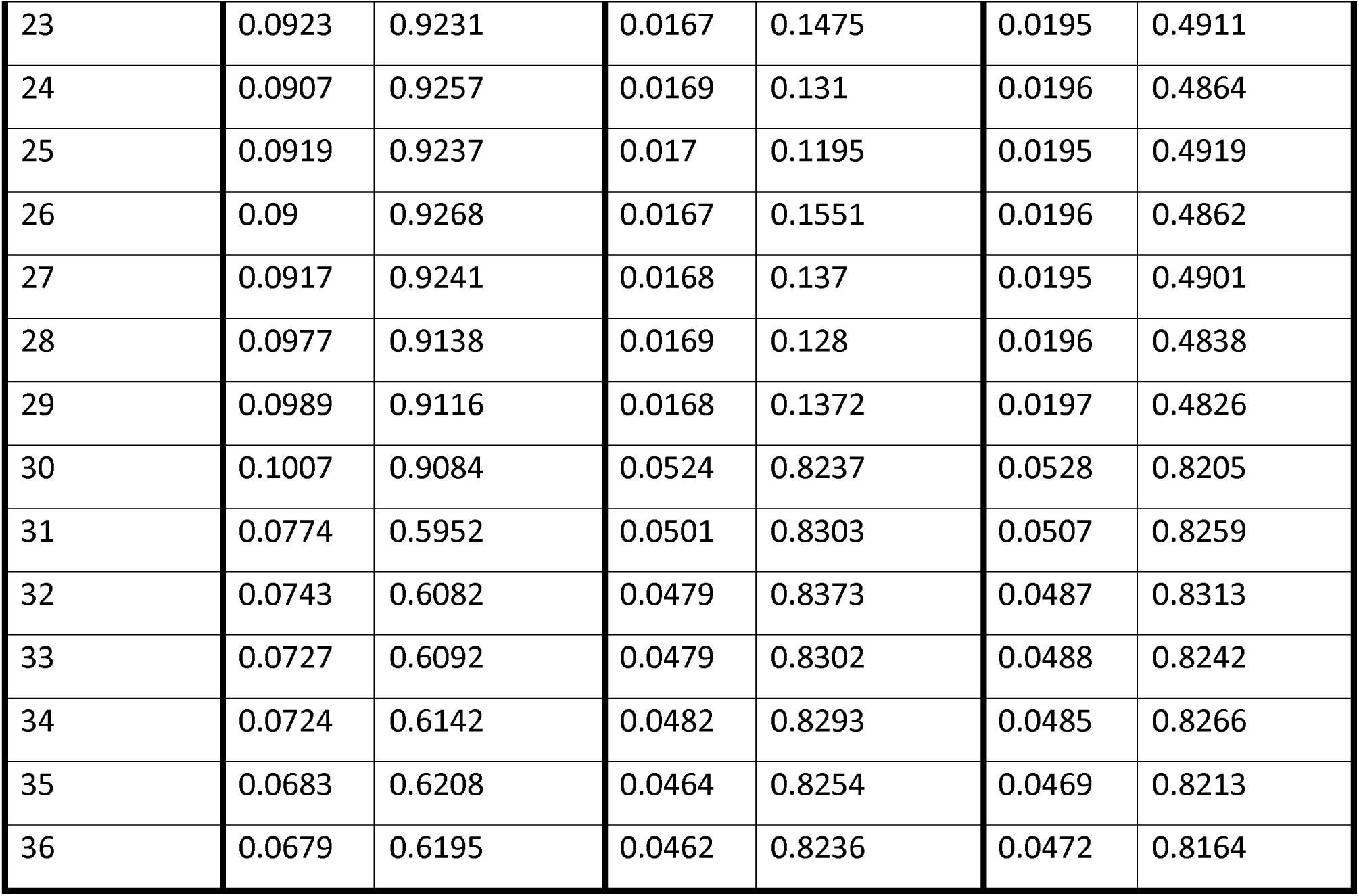
Third-degree polynomial fit. Correlations between MSE and *dynamic* nodal network features

**Table 4. 8.**
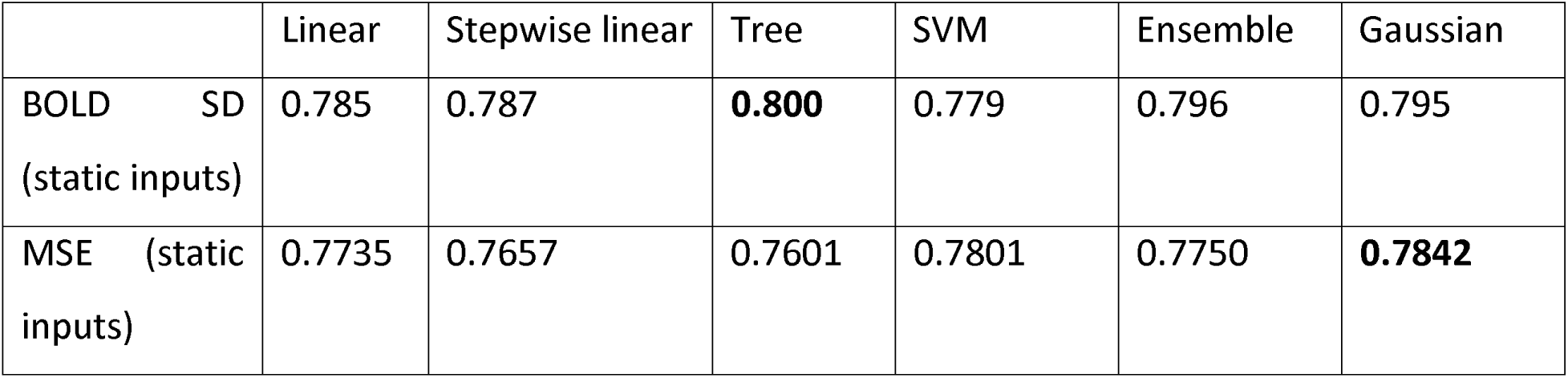
Accuracy of MSE and BOLD SD predictions based on static nodal brain network measures using different prediction models. Accuracy values were averaged over all brain regions.

|  | Linear | Stepwise linear | Tree | SVM | Ensemble | Gaussian |
| --- | --- | --- | --- | --- | --- | --- |
| BOLD SD<br>(static inputs) | 0.785 | 0.787 | **0.800** | 0.779 | 0.796 | 0.795 |
| MSE (static inputs) | 0.7735 | 0.7657 | 0.7601 | 0.7801 | 0.7750 | **0.7842** |

Prediction accuracy for MSE scales using six different linear and nonlinear approaches is shown in Figure 4.8. Similarly to SD prediction, for each approach, inputs were the static nodal measures (i.e., BC, EC, CC), and the outputs were regional MSE at different scales, in separate models. In general, all approaches showed reliable predictions for MSE at fine scales (scales 1 and 2) (average accuracy over all regions and models was 0.94). However, we observed a reduction in prediction to an average accuracy of 0.76 for coarse scales (scales 3 to 36) particularly in the temporal pole, inferior temporal and olfactory cortices, hippocampus and parahippocampal, amygdala, pallidum, and thalamus. In general, the best MSE prediction result was obtained using the Gaussian approach 0.78 (see Table 4.8).

### 3-5 Prediction of SD and MSE using dynamical nodal network measures

We used six models to predict SD using the Shannon entropy of nodal measures (i.e., BC, EC, CC) as inputs. The average accuracy of SD prediction across all models and all regions was reliable (accuracy>0.78), and the *tree* model yielded the highest accuracy over all regions with an average value of 0.80 (see Figure 4.9 and Table 4.9).

**Figure 4. 9.**
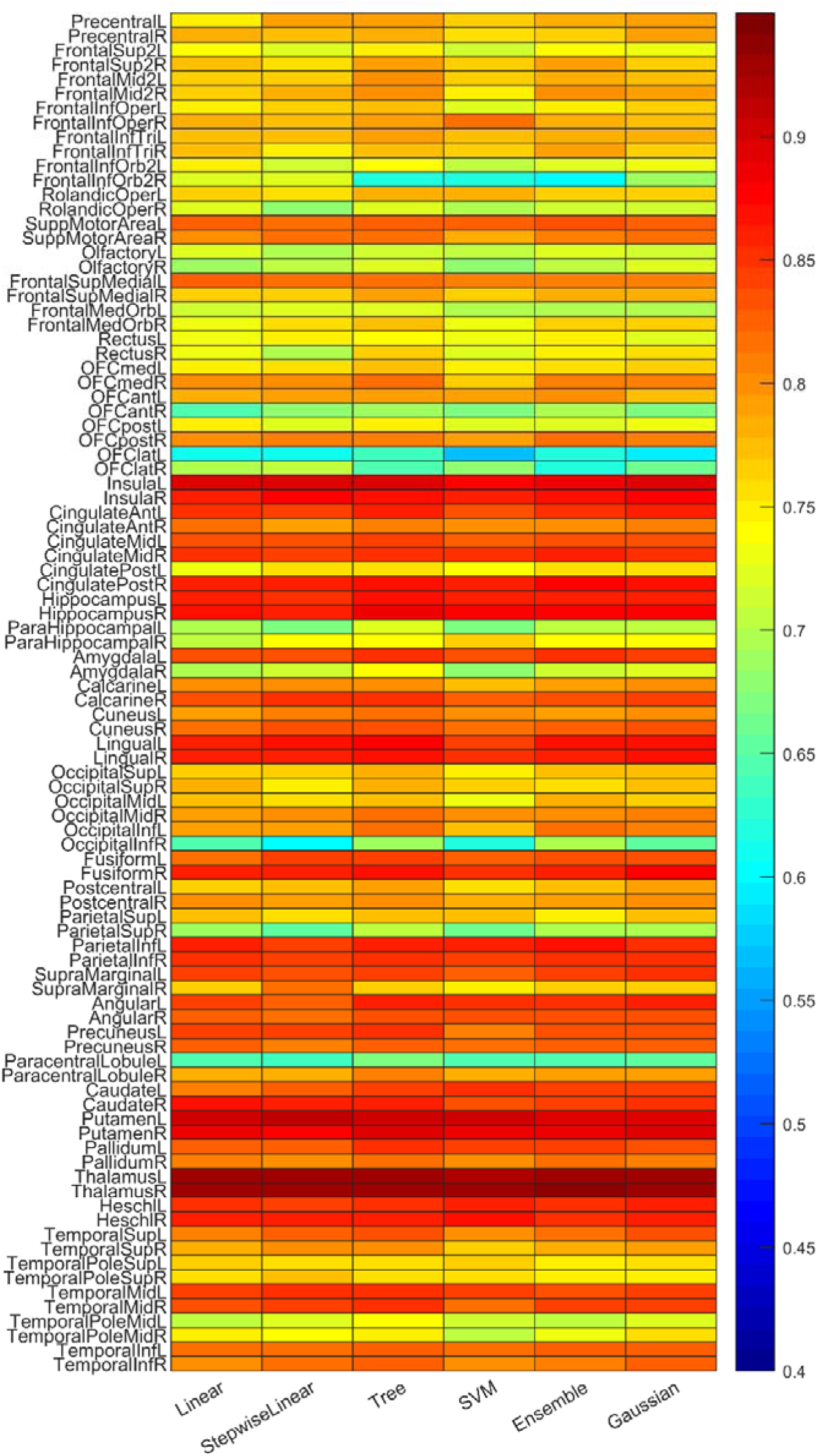
BOLD SD prediction accuracy based on *dynamic* nodal brain network features (BC, EC, CC).

**Table 4. 9.** Accuracy of MSE and BOLD SD predictions based on dynamic nodal brain network measures using different prediction models. Accuracy values were averaged over all brain regions.

|  | Linear | Stepwise linear | Tree | SVM | Ensemble | Gaussian |
| --- | --- | --- | --- | --- | --- | --- |
| BOLD SD<br>(dynamic inputs) | 0.790 | 0.788 | **0.801** | 0.781 | 0.792 | 0.794 |
| MSE-fine scale<br>(dynamic inputs) | **0.7854** | 0.7801 | 0.7600 | 0.7779 | 0.7751 | 0.7847 |

The accuracy of the six models for MSE prediction at fine scales was high with an average value of 0.95. Similarly to our statics results, MSE prediction accuracy was reduced to an average of 0.77 for coarse scales, especially in the temporal pole, inferior temporal and olfactory cortices, hippocampus and parahippocampal, amygdala, pallidum, and thalamus. In general, the linear model yielded the best prediction for MSE with an average accuracy value of 0.79 (see Figure 4.10 and Table 4.9).

**Figure 4. 10.**
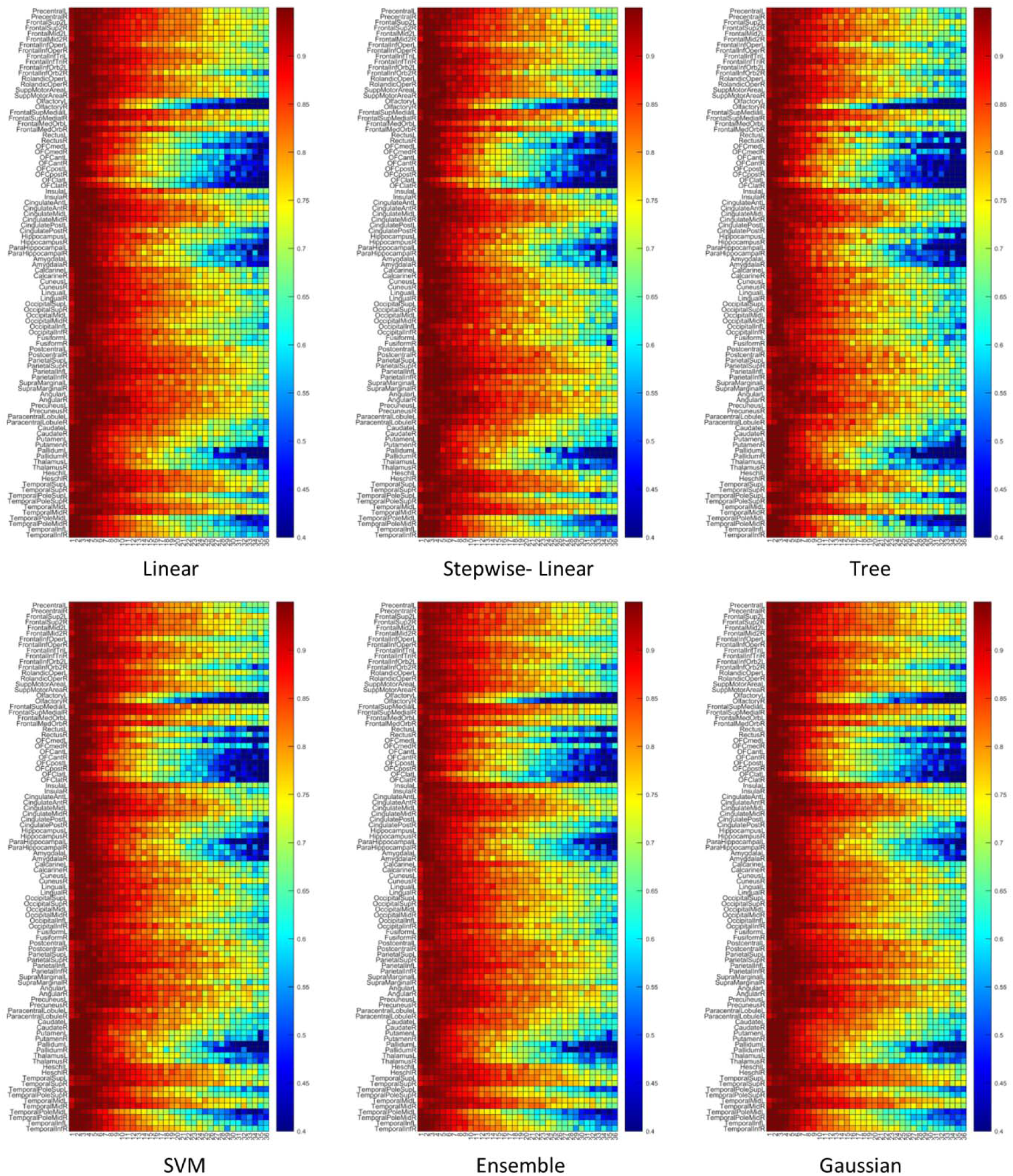
MSE prediction accuracy based on *dynamic* nodal network features (BC, EC, CC).

## 4 Discussion

We examined the relationship between BOLD variability, complexity, static/dynamic functional brain network features (represented by nodal measures from graph theory analysis) for healthy participants during movie watching. We found that the correlation between brain signal variability (quantified using BOLD SD) and brain signal complexity (quantified using BOLD MSE) was dependent on time scale of MSE, with a positive relationship at fine scales but a negative relationship at coarser scales. In addition, regions with high centrality and high clustering coefficient were associated with lower variability, as well as lower complexity at fine time scales. This relationship was reversed at coarser time scales. A similar relationship existed for dynamic measures, but contrary to our hypotheses, this relationship was weaker, as it was significant only for some aspects of regional centrality dynamics (i.e., entropy of BC). Prediction results from linear and nonlinear models confirmed the relationships between nodal network measures with BOLD SD and MSE. It also suggested that nodal connectivity patterns in functional brain network can demonstrate the fluctuations and complexity of BOLD signals in brain regions. These findings highlight the interplay between metrics that index the dynamics and complexity of brain signal, as well as how they link to static and dynamic functional brain network metrics in healthy adults.

### 4-1 The relationship between BOLD SD and MSE

Brain signal variability and complexity are both dynamic measures that reflect the adaptive capability of the brain (Goldberger, Peng, & Lipsitz, 2002; Grady & Garrett, 2018). Although these two measures are correlated, they are not equivalent (Lipsitz, 2002; Van Emmerik et al., 2016). The current study revealed a positive correlation between BOLD SD with MSE at the finest time scales (1 to 2 second windows), and a negative correlation at more coarse time scales (3 to 36 second windows). One possible reason that BOLD SD and MSE were positively correlated at the finest scale is that they both were computed from the original time series. MSE estimation is based on the similarity criterion r, which is a proportion of the standard deviation of the original time series. Starting from the second scale, the time series for MSE calculation is shorter and smoother due to the nature of coarse graining process, which acts as a low pass filter. In other words, the standard deviation of the new time series becomes smaller. but the similarity criterion remains unchanged, resulting in smaller MSE values at the second scale (Kosciessa, Kloosterman, & Garrett, 2020). As MSE scale increases, the SD of the new time series decreases and MSE values also decrease. Therefore, the correlation between SD estimated from the original time series and MSE at coarser time scales becomes negative.

### 4-2 BOLD SD and MSE relate to static nodal functional brain network features

Using regression analysis, we observed a negative correlation between BOLD SD and nodal measures (i.e., BC, EC and CC). In other words, a brain network hub (i.e., regions with high BC and EC) or a highly clustered region (i.e., regions with high CC) is associated with less variable signal. Theoretical and empirical work has linked lower brain signal variability to better signal transmission (He, 2013; Ponce-Alvarez, He, Hagmann, & Deco, 2015). Thus, we propose that lower regional signal variability may be related to consistency in connectivity patterns in hubs and highly clustered regions, for better information processing. A recent fMRI study may support this argument by demonstrating that in both resting and task states, nodes with lower CC change their memberships across different modules frequently (Yi, Fan, & Wu, 2022). Therefore, less regional signal variability for a highly clustered region (node with greater CC) may indicate that the membership of the region in a certain network is consistent.

Our regression analyses additionally revealed a negative correlation between nodal measures (i.e., BC, EC and CC) with MSE at fine scales (i.e., the first and second scales), but a positive correlation with MSE at coarser scales. This finding is consistent with a previous fMRI study that showed an association between functional connectivity and BOLD MSE, which was negative at fine scales and positive at coarse scales in resting state networks (McDonough & Nashiro, 2014). The authors argued that their findings are consistent with a neural model of information processing (Baptista & Kurths, 2008), which suggests that information processing capacity was higher (comparable to greater MSE) when a network showed greater synchrony (comparable to greater functional connectivity) at coarse time scales and greater desynchrony (lower functional connectivity) at fine time scales (McDonough & Nashiro, 2014).

The prediction results of the linear and nonlinear models also confirm the relationships between nodal features in functional brain network and BOLD SD and MSE. In particular, prediction accuracy of fine scale MSE based on static nodal network features was higher than that for coarse scale MSE. Previous work suggests that fine scale MSE is related to local neural processing and coarse scale MSE represents long-distance communication between regions (McDonough & Nashiro, 2014; McIntosh et al., 2014). Since the predictors (BC, EC, and CC) were of nodal functional network features, it is reasonable that local information processing represented by fine scale MSE is better predicted.

### 4-3 BOLD SD and MSE relate to dynamic nodal functional brain network features

Although we observed significant correlations between SD and MSE with dynamic functional brain network measures, contrary to our expectations, these relationships were not stronger for dynamic as compared to static metrics. Specifically, we observed a negative correlation between BOLD SD and the Shannon entropy of nodal measures across time. However, correlations were significant only for BC and CC, but not EC. Similarly, we found a negative correlation between the Shannon entropy of nodal measures across time with MSE at fine scales, and a positive correlation with MSE at coarser time scales. Again, these correlations were significant only for BC and CC. These findings first revealed links between SD and MSE with the dynamics of a hubness measure (i.e., entropy of nodal BC) and a nodal segregation measure (i.e., entropy of nodal CC) which is similar to the association for static nodal network features. But we also found no correlation with another measure of hubness (i.e., entropy of nodal EC). One potential explanation is that the reorganization of functional networks across time is a complex scenario. Previous theoretical work using surrogate dynamic functional connectivity models and random walk analyses has shown that functional connectivity fluctuation over time does not simply maintain static functional connectivity features, nor does it show fully unrelated patterns (Battaglia et al., 2020). Instead, time-varying functional network reconfigurations demonstrate long-range sequential correlations which are indicative of a complex progress that is neither completely orderly nor completely random (Battaglia et al., 2020; Crutchfield, 2012). Given that dynamic functional connectivity and brain networks are intrinsically complex, we argue that the associations between SD and MSE with functional brain network measures vary with time in a complex way. Specifically, it keeps certain features of the static functional brain networks on the one hand, e.g., the relationship between SD and MSE with some aspects of centrality and clustering coefficient of regions; but introduces some degree of deviation on the other hand, e.g., a nonsignificant relationship between SD and MSE with other aspects of hubness.

One may wonder why the dynamic association disappears in *EC* but not in *BC*. One possible reason is the local dependence of *EC* on the degree of a node and its neighbors (Ruhnau, 2000). In short time scales, the degree of a node can be highly influenced by alterations in regional connectivity resulting in a high level of degree uncertainty. This increased uncertainty can increase nonlinearity in the dynamics of *EC* (which is directly associated with degree) and also increase the entropy of *EC,* resulting in a ceiling effect for Shannon entropy values, and a reduced correlation with SD and MSE. *BC*, on the other hand, is associated with shortest paths in the networks (Boccaletti et al., 2006) and are possibly less affected by regional alterations at short time scales. As we show in Figure 4, the Shannon entropy of *BC* exhibited sufficient heterogeneity in values across regions to allow them to be related to the complexity of activity in different regions.

## 5 Limitations

The present study has several limitations. First, clustering coefficients were calculated from a correlation matrix where each entity of the correlation matrix was a correlation coefficient representing functional connectivity between two corresponding ROIs. Previous work suggests that a network generated by a correlation matrix is likely to have more indirect paths which increases the values of the clustering coefficient (Adachi et al., 2012; Zalesky, Fornito, & Bullmore, 2012). Future studies should examine clustering coefficients calculated from partial mutual information or three-way partial correlation coefficients to reduce the potential influence of indirect paths (Masuda, Sakaki, Ezaki, & Watanabe, 2018). Second, data in the current study was acquired from 1.5 T scanner. We used this dataset because we were interested in comparing signal variability and complexity directly, and the extended length of functional acquisition in this dataset made it ideal for estimating brain signal complexity (MSE) over multiple time scales. Given that the SNR is approximately 25% lower for 1.5T as compared to 3T acquisition (Wardlaw et al., 2012), replication using data collected at higher magnetic field strength is essential. Lastly, although the age range of participants is 18-55 years, we did not test for age effects because only four participants were over 35 years of age. Future work should examine whether or not the relationship between measures of variability, complexity, and static and dynamic functional connectivity differs with age.

## 6 Conclusions

The current study examined brain signal variability (i.e., BOLD SD) and complexity (i.e., BOLD MSE) simultaneously to elucidate differences in their relationship with static and dynamic measures of functional brain networks. Our findings suggest that the relationship between SD and MSE is dependent on the temporal scale of MSE, with a positive correlation at fine scales and a negative correlation at coarse scales. Consequently, the two metrics correlated with static and dynamic functional brain network measures in opposite directions at coarse MSE scales. Specifically, regions with high centrality and clustering coefficient were related to less variable (lower BOLD SD) but more complex (greater coarse scale MSE) signal. A similar relationship existed for dynamic measures, but this relationship was weaker, as it was not significant for some aspects of regional centrality dynamics (i.e., entropy of EC). These findings may be related to the complex, time-varying feature of functional brain networks and reflect how BOLD SD and MSE co-evolve with dynamic functional brain networks over time.

## Notes

### Competing Interest Statement

The authors have declared no competing interest.

### Summary of Updates

A few edits have been applied.

