## Supplementary Tables for "Exploring the Interplay between BOLD Signal Variability, Complexity, and Static and Dynamic Functional Brain Network Features during Movie Viewing"

Table S1

Linear regression: Correlations between MSE and *static* nodal network features

| | **Betweenness Centrality (BC)** | | | | **Eigenvector Centrality (EC)** | | | | **Clustering Coefficient (CC)** | | | |
|---|---|---|---|---|---|---|---|---|---|---|---|---|
| MSE time scale | RMSE | R-squared | t-statistic | p-value | RMSE | R-squared | t-statistic | p-value | RMSE | R-squared | t-statistic | p-value |
| 1 | 0.0823 | 0.4354 | -8.4226 | <1e-5 | 0.0574 | 0.725 | -15.5756 | <1e-10 | 0.0619 | 0.6799 | -13.9804 | <1e-09 |
| 2 | 0.0256 | 0.4024 | -7.8701 | 1.276e-5 | 0.0266 | 0.3586 | -7.1712 | 1.833e-10 | 0.0273 | 0.3231 | -6.6272 | 2.282e-09 |
| 3 | 0.0386 | 0.1879 | 4.6137 | <1e-5 | 0.0253 | 0.6515 | 13.1136 | <1e-10 | 0.0257 | 0.64 | 12.7896 | <1e-09 |
| 4 | 0.0645 | 0.3236 | 6.6347 | <1e-5 | 0.0372 | 0.7747 | 17.7864 | <1e-10 | 0.0389 | 0.7545 | 16.8144 | <1e-09 |
| 5 | 0.081 | 0.355 | 7.1155 | <1e-5 | 0.0467 | 0.7852 | 18.339 | <1e-10 | 0.0489 | 0.7646 | 17.2878 | <1e-09 |
| 6 | 0.0907 | 0.387 | 7.6212 | <1e-5 | 0.0522 | 0.7969 | 18.9977 | <1e-10 | 0.0547 | 0.777 | 17.904 | <1e-09 |
| 7 | 0.0971 | 0.4105 | 8.0041 | <1e-5 | 0.0566 | 0.7998 | 19.1715 | <1e-10 | 0.0591 | 0.7819 | 18.1632 | <1e-09 |
| 8 | 0.1003 | 0.4328 | 8.3787 | <1e-5 | 0.0587 | 0.8058 | 19.5393 | <1e-10 | 0.0615 | 0.7871 | 18.4414 | <1e-09 |
| 9 | 0.104 | 0.4395 | 8.4938 | <1e-5 | 0.0611 | 0.8063 | 19.571 | <1e-10 | 0.0641 | 0.7869 | 18.429 | <1e-09 |
| 10 | 0.1056 | 0.4589 | 8.8323 | <1e-5 | 0.0609 | 0.82 | 20.4704 | <1e-10 | 0.0641 | 0.8009 | 19.2395 | <1e-09 |
| 11 | 0.1075 | 0.4704 | 9.0401 | <1e-5 | 0.0623 | 0.822 | 20.6116 | <1e-10 | 0.0655 | 0.8035 | 19.3986 | <1e-09 |
| 12 | 0.1077 | 0.4842 | 9.2939 | <1e-5 | 0.0628 | 0.8244 | 20.7844 | <1e-10 | 0.066 | 0.8059 | 19.5446 | <1e-09 |
| 13 | 0.1072 | 0.4964 | 9.5237 | <1e-5 | 0.0625 | 0.8289 | 21.1115 | <1e-10 | 0.0655 | 0.8124 | 19.9603 | <1e-09 |
| 14 | 0.1079 | 0.5027 | 9.6445 | <1e-5 | 0.0641 | 0.8246 | 20.7977 | <1e-10 | 0.0671 | 0.8079 | 19.6687 | <1e-09 |
| 15 | 0.1072 | 0.511 | 9.8059 | <1e-5 | 0.0633 | 0.8292 | 21.1302 | <1e-10 | 0.0662 | 0.8136 | 20.0394 | <1e-09 |
| 16 | 0.1049 | 0.5284 | 10.1536 | <1e-5 | 0.062 | 0.8356 | 21.6239 | <1e-10 | 0.0648 | 0.8203 | 20.4901 | <1e-09 |
| 17 | 0.1049 | 0.526 | 10.1049 | <1e-5 | 0.0623 | 0.8329 | 21.4166 | <1e-10 | 0.0646 | 0.82 | 20.4705 | <1e-09 |
| 18 | 0.103 | 0.5453 | 10.5037 | <1e-5 | 0.0613 | 0.839 | 21.8958 | <1e-10 | 0.0638 | 0.8255 | 20.8631 | <1e-09 |
| 19 | 0.1012 | 0.544 | 10.4755 | <1e-5 | 0.0608 | 0.8355 | 21.6138 | <1e-10 | 0.0632 | 0.8224 | 20.6369 | <1e-09 |
| 20 | 0.1 | 0.5496 | 10.5949 | <1e-5 | 0.0595 | 0.8405 | 22.0208 | <1e-10 | 0.0619 | 0.8272 | 20.9881 | <1e-09 |
| 21 | 0.097 | 0.5616 | 10.8561 | <1e-5 | 0.0595 | 0.8348 | 21.5621 | <1e-10 | 0.0612 | 0.8255 | 20.8638 | <1e-09 |
| 22 | 0.0968 | 0.5621 | 10.8677 | <1e-5 | 0.059 | 0.8372 | 21.7512 | <1e-10 | 0.0606 | 0.8284 | 21.0772 | <1e-09 |
| 23 | 0.0935 | 0.5708 | 11.0622 | <1e-5 | 0.0591 | 0.8285 | 21.0791 | <1e-10 | 0.061 | 0.8174 | 20.2952 | <1e-09 |
| 24 | 0.092 | 0.5779 | 11.2223 | <1e-5 | 0.0582 | 0.831 | 21.2723 | <1e-10 | 0.0599 | 0.8211 | 20.5512 | <1e-09 |
| 25 | 0.0884 | 0.5848 | 11.3844 | <1e-5 | 0.0563 | 0.8318 | 21.3288 | <1e-10 | 0.0575 | 0.8241 | 20.7634 | <1e-09 |
| 26 | 0.0888 | 0.5825 | 11.3307 | <1e-5 | 0.0538 | 0.8469 | 22.5575 | <1e-10 | 0.0557 | 0.8362 | 21.6686 | <1e-09 |
| 27 | 0.0869 | 0.5766 | 11.1933 | <1e-5 | 0.0536 | 0.839 | 21.895 | <1e-10 | 0.0549 | 0.8308 | 21.2515 | <1e-09 |
| 28 | 0.0847 | 0.5801 | 11.2728 | <1e-5 | 0.0545 | 0.8262 | 20.9151 | <1e-10 | 0.0556 | 0.8188 | 20.3919 | <1e-09 |

| | | | | | | | | | | | | |
|---|---|---|---|---|---|---|---|---|---|---|---|---|
| 29 | 0.0814 | 0.5924 | 11.564 | <1e-5 | 0.0531 | 0.8262 | 20.9147 | <1e-10 | 0.0541 | 0.82 | 20.4753 | <1e-09 |
| 30 | 0.079 | 0.5985 | 11.71 | <1e-5 | 0.0524 | 0.8237 | 20.7315 | <1e-10 | 0.0528 | 0.8205 | 20.507 | <1e-09 |
| 31 | 0.0774 | 0.5952 | 11.6304 | <1e-5 | 0.0501 | 0.8303 | 21.2159 | <1e-10 | 0.0507 | 0.8259 | 20.888 | <1e-09 |
| 32 | 0.0743 | 0.6082 | 11.9515 | <1e-5 | 0.0479 | 0.8373 | 21.7606 | <1e-10 | 0.0487 | 0.8313 | 21.2954 | <1e-09 |
| 33 | 0.0727 | 0.6092 | 11.975 | <1e-5 | 0.0479 | 0.8302 | 21.2115 | <1e-10 | 0.0488 | 0.8242 | 20.7669 | <1e-09 |
| 34 | 0.0724 | 0.6142 | 12.1034 | <1e-5 | 0.0482 | 0.8293 | 21.1448 | <1e-10 | 0.0485 | 0.8266 | 20.9449 | <1e-09 |
| 35 | 0.0683 | 0.6208 | 12.2722 | <1e-5 | 0.0464 | 0.8254 | 20.8573 | <1e-10 | 0.0469 | 0.8213 | 20.5625 | <1e-09 |
| 36 | 0.0679 | 0.6195 | 12.2379 | <1e-5 | 0.0462 | 0.8236 | 20.7255 | <1e-10 | 0.0472 | 0.8164 | 20.2238 | <1e-09 |

Table S2
Third-degree polynomial fit: Correlations between MSE and *static* nodal network features

| | **Betweenness Centrality (BC)** | | **Eigenvector Centrality (EC)** | | **Clustering Coefficient (CC)** | |
|---|---|---|---|---|---|---|
| MSE time scale | RMSE | R-squared | RMSE | R-squared | RMSE | R-squared |
| 1 | 0.0823 | 0.4354 | 0.0574 | 0.725 | 0.0619 | 0.6799 |
| 2 | 0.0256 | 0.4024 | 0.0266 | 0.3586 | 0.0273 | 0.3231 |
| 3 | 0.0386 | 0.1879 | 0.0253 | 0.6515 | 0.0257 | 0.64 |
| 4 | 0.0645 | 0.3236 | 0.0372 | 0.7747 | 0.0389 | 0.7545 |
| 5 | 0.081 | 0.355 | 0.0467 | 0.7852 | 0.0489 | 0.7646 |
| 6 | 0.0907 | 0.387 | 0.0522 | 0.7969 | 0.0547 | 0.777 |
| 7 | 0.0971 | 0.4105 | 0.0566 | 0.7998 | 0.0591 | 0.7819 |
| 8 | 0.1003 | 0.4328 | 0.0587 | 0.8058 | 0.0615 | 0.7871 |
| 9 | 0.104 | 0.4395 | 0.0611 | 0.8063 | 0.0641 | 0.7869 |
| 10 | 0.1056 | 0.4589 | 0.0609 | 0.82 | 0.0641 | 0.8009 |
| 11 | 0.1075 | 0.4704 | 0.0623 | 0.822 | 0.0655 | 0.8035 |
| 12 | 0.1077 | 0.4842 | 0.0628 | 0.8244 | 0.066 | 0.8059 |
| 13 | 0.1072 | 0.4964 | 0.0625 | 0.8289 | 0.0655 | 0.8124 |
| 14 | 0.1079 | 0.5027 | 0.0641 | 0.8246 | 0.0671 | 0.8079 |
| 15 | 0.1072 | 0.511 | 0.0633 | 0.8292 | 0.0662 | 0.8136 |
| 16 | 0.1049 | 0.5284 | 0.062 | 0.8356 | 0.0648 | 0.8203 |
| 17 | 0.1049 | 0.526 | 0.0623 | 0.8329 | 0.0646 | 0.82 |
| 18 | 0.103 | 0.5453 | 0.0613 | 0.839 | 0.0638 | 0.8255 |
| 19 | 0.1012 | 0.544 | 0.0608 | 0.8355 | 0.0632 | 0.8224 |
| 20 | 0.1 | 0.5496 | 0.0595 | 0.8405 | 0.0619 | 0.8272 |
| 21 | 0.097 | 0.5616 | 0.0595 | 0.8348 | 0.0612 | 0.8255 |
| 22 | 0.0968 | 0.5621 | 0.059 | 0.8372 | 0.0606 | 0.8284 |
| 23 | 0.0935 | 0.5708 | 0.0591 | 0.8285 | 0.061 | 0.8174 |
| 24 | 0.092 | 0.5779 | 0.0582 | 0.831 | 0.0599 | 0.8211 |
| 25 | 0.0884 | 0.5848 | 0.0563 | 0.8318 | 0.0575 | 0.8241 |
| 26 | 0.0888 | 0.5825 | 0.0538 | 0.8469 | 0.0557 | 0.8362 |
| 27 | 0.0869 | 0.5766 | 0.0536 | 0.839 | 0.0549 | 0.8308 |
| 28 | 0.0847 | 0.5801 | 0.0545 | 0.8262 | 0.0556 | 0.8188 |
| 29 | 0.0814 | 0.5924 | 0.0531 | 0.8262 | 0.0541 | 0.82 |
| 30 | 0.079 | 0.5985 | 0.0524 | 0.8237 | 0.0528 | 0.8205 |

| | | | | | | |
|---|---|---|---|---|---|---|
| 31 | 0.0774 | 0.5952 | 0.0501 | 0.8303 | 0.0507 | 0.8259 |
| 32 | 0.0743 | 0.6082 | 0.0479 | 0.8373 | 0.0487 | 0.8313 |
| 33 | 0.0727 | 0.6092 | 0.0479 | 0.8302 | 0.0488 | 0.8242 |
| 34 | 0.0724 | 0.6142 | 0.0482 | 0.8293 | 0.0485 | 0.8266 |
| 35 | 0.0683 | 0.6208 | 0.0464 | 0.8254 | 0.0469 | 0.8213 |
| 36 | 0.0679 | 0.6195 | 0.0462 | 0.8236 | 0.0472 | 0.8164 |

Table S3
Linear regression: Correlations between MSE and *dynamic* nodal network features

| | **Dynamic Betweenness Centrality (dBC)** | | | | **Dynamic Eigenvector Centrality (dEC)** | | | | **Dynamic Clustering Coefficient (dCC)** | | | |
|---|---|---|---|---|---|---|---|---|---|---|---|---|
| MSE time scale | RMSE | R-squared | t-statistic | p-value | RMSE | R-squared | t-statistic | p-value | RMSE | R-squared | t-statistic | p-value |
| 1 | 0.1129 | 0.8875 | -26.9425 | <1e-13 | 0.0183 | 0.0058 | 0.7306 | 0.4669 | 0.0231 | 0.2998 | -6.2769 | <1e-03 |
| 2 | 0.2497 | 0.4494 | -8.6659 | 1.461e-13 | 0.0179 | 0.0499 | 2.1973 | 0.0305 | 0.0266 | 0.0747 | -2.7261 | 7.2e-03 |
| 3 | 0.1576 | 0.7805 | 18.0874 | <1e-13 | 0.0182 | 0.0189 | 1.3298 | 0.1869 | 0.0205 | 0.4497 | 8.6708 | <1e-03 |
| 4 | 0.0853 | 0.9358 | 36.6121 | <1e-13 | 0.0183 | 0.0037 | 0.5843 | 0.5604 | 0.0205 | 0.4498 | 8.6724 | <1e-03 |
| 5 | 0.0712 | 0.9552 | 44.3129 | <1e-13 | 0.0183 | 0.0023 | 0.4557 | 0.6497 | 0.0205 | 0.4498 | 8.6729 | <1e-03 |
| 6 | 0.0609 | 0.9673 | 52.1379 | <1e-13 | 0.0183 | 0.0011 | 0.3252 | 0.7457 | 0.0204 | 0.4533 | 8.7337 | <1e-03 |
| 7 | 0.0592 | 0.9691 | 53.6801 | <1e-13 | 0.0183 | 0.0006 | 0.2314 | 0.8175 | 0.0206 | 0.4429 | 8.5524 | <1e-03 |
| 8 | 0.0605 | 0.9676 | 52.4449 | <1e-13 | 0.0183 | 0.0003 | 0.173 | 0.863 | 0.0204 | 0.455 | 8.7632 | <1e-03 |
| 9 | 0.0614 | 0.9667 | 51.6411 | <1e-13 | 0.0183 | 0 | 0.032 | 0.9745 | 0.0206 | 0.4465 | 8.6153 | <1e-03 |
| 10 | 0.0616 | 0.9665 | 51.5272 | <1e-13 | 0.0183 | 0 | -0.0269 | 0.9786 | 0.0204 | 0.4551 | 8.7653 | <1e-03 |
| 11 | 0.0628 | 0.9651 | 50.4756 | <1e-13 | 0.0183 | 0.0002 | -0.1252 | 0.9007 | 0.0204 | 0.4574 | 8.8071 | <1e-03 |
| 12 | 0.0677 | 0.9596 | 46.7302 | <1e-13 | 0.0183 | 0.0004 | -0.1891 | 0.8504 | 0.0202 | 0.4638 | 8.9202 | <1e-03 |
| 13 | 0.0701 | 0.9566 | 45.0365 | <1e-13 | 0.0183 | 0.0006 | -0.226 | 0.8217 | 0.0203 | 0.4602 | 8.8567 | <1e-03 |
| 14 | 0.0749 | 0.9505 | 42.0293 | <1e-13 | 0.0183 | 0.0009 | -0.2838 | 0.7772 | 0.0203 | 0.4612 | 8.8747 | <1e-03 |
| 15 | 0.0769 | 0.9478 | 40.8582 | <1e-13 | 0.0183 | 0.0015 | -0.3707 | 0.7117 | 0.0202 | 0.4651 | 8.9449 | <1e-03 |
| 16 | 0.08 | 0.9435 | 39.192 | <1e-13 | 0.0183 | 0.0024 | -0.4744 | 0.6364 | 0.0202 | 0.4637 | 8.9184 | <1e-03 |
| 17 | 0.082 | 0.9407 | 38.1867 | <1e-13 | 0.0183 | 0.0017 | -0.4007 | 0.6896 | 0.0203 | 0.4611 | 8.8718 | <1e-03 |
| 18 | 0.0865 | 0.9339 | 36.062 | <1e-13 | 0.0183 | 0.0032 | -0.5408 | 0.5899 | 0.0201 | 0.4688 | 9.011 | <1e-03 |
| 19 | 0.0899 | 0.9286 | 34.5892 | <1e-13 | 0.0183 | 0.0027 | -0.5003 | 0.618 | 0.0202 | 0.4642 | 8.9286 | <1e-03 |
| 20 | 0.0943 | 0.9215 | 32.8524 | <1e-13 | 0.0183 | 0.0042 | -0.6232 | 0.5347 | 0.0202 | 0.4674 | 8.9846 | <1e-03 |
| 21 | 0.0969 | 0.9171 | 31.8944 | <1e-13 | 0.0183 | 0.0041 | -0.6161 | 0.5394 | 0.02 | 0.4749 | 9.1215 | <1e-03 |
| 22 | 0.0965 | 0.9178 | 32.0502 | <1e-13 | 0.0183 | 0.0037 | -0.5878 | 0.5581 | 0.0202 | 0.4678 | 8.9931 | <1e-03 |
| 23 | 0.1063 | 0.9002 | 28.8077 | <1e-13 | 0.0183 | 0.0057 | -0.7281 | 0.4684 | 0.0201 | 0.4717 | 9.0626 | <1e-03 |
| 24 | 0.1071 | 0.8987 | 28.566 | <1e-13 | 0.0183 | 0.0063 | -0.7667 | 0.4452 | 0.0201 | 0.4696 | 9.0256 | <1e-03 |
| 25 | 0.1095 | 0.8941 | 27.8629 | <1e-13 | 0.0183 | 0.005 | -0.6802 | 0.4981 | 0.0199 | 0.4793 | 9.2033 | <1e-03 |
| 26 | 0.1064 | 0.8999 | 28.7666 | <1e-13 | 0.0183 | 0.0071 | -0.8113 | 0.4193 | 0.0201 | 0.4707 | 9.0454 | <1e-03 |
| 27 | 0.108 | 0.897 | 28.3014 | <1e-13 | 0.0183 | 0.006 | -0.7425 | 0.4597 | 0.0201 | 0.4731 | 9.0884 | <1e-03 |
| 28 | 0.1129 | 0.8875 | 26.9337 | <1e-13 | 0.0183 | 0.0069 | -0.8019 | 0.4247 | 0.0201 | 0.4692 | 9.0175 | <1e-03 |

| | | | | | | | | | | | | |
|---|---|---|---|---|---|---|---|---|---|---|---|---|
| 29 | 0.1166 | 0.8799 | 25.9625 | <1e-13 | 0.0183 | 0.008 | -0.8614 | 0.3912 | 0.0202 | 0.4672 | 8.9821 | <1e-03 |
| 30 | 0.1194 | 0.874 | 25.2653 | <1e-13 | 0.0183 | 0.0081 | -0.8678 | 0.3878 | 0.0202 | 0.4665 | 8.9692 | <1e-03 |
| 31 | 0.1177 | 0.8776 | 25.6796 | <1e-13 | 0.0182 | 0.0094 | -0.9351 | 0.3522 | 0.0202 | 0.4648 | 8.9392 | <1e-03 |
| 32 | 0.1201 | 0.8726 | 25.1061 | <1e-13 | 0.0182 | 0.009 | -0.9125 | 0.3639 | 0.0201 | 0.4718 | 9.0658 | <1e-03 |
| 33 | 0.1205 | 0.8718 | 25.0145 | <1e-13 | 0.0182 | 0.0092 | -0.9252 | 0.3573 | 0.0202 | 0.4642 | 8.9279 | <1e-03 |
| 34 | 0.1243 | 0.8635 | 24.1263 | <1e-13 | 0.0183 | 0.0087 | -0.8961 | 0.3725 | 0.0201 | 0.4727 | 9.0808 | <1e-03 |
| 35 | 0.129 | 0.8531 | 23.1132 | <1e-13 | 0.0182 | 0.0111 | -1.014 | 0.3132 | 0.0203 | 0.4595 | 8.8433 | <1e-03 |
| 36 | 0.1345 | 0.8401 | 21.9893 | <1e-13 | 0.0182 | 0.0142 | -1.1515 | 0.2525 | 0.0203 | 0.4633 | 8.911 | <1e-03 |

Table S4

Third-degree polynomial fit: Correlations between MSE and *dynamic* nodal network features

| | **Dynamic Betweenness Centrality (dBC)** | | **Dynamic Eigenvector Centrality (dEC)** | | **Dynamic Clustering Coefficient (dCC)** | |
|---|---|---|---|---|---|---|
| MSE time scale | RMSE | R-squared | RMSE | R-squared | RMSE | R-squared |
| 1 | 0.0614 | 0.9659 | 0.0168 | 0.1424 | 0.0196 | 0.4884 |
| 2 | 0.2294 | 0.525 | 0.0173 | 0.0862 | 0.0252 | 0.1526 |
| 3 | 0.1437 | 0.8136 | 0.0177 | 0.0443 | 0.0195 | 0.4902 |
| 4 | 0.0753 | 0.9488 | 0.0173 | 0.0868 | 0.0194 | 0.4988 |
| 5 | 0.0629 | 0.9643 | 0.0173 | 0.0947 | 0.019 | 0.5181 |
| 6 | 0.0541 | 0.9736 | 0.0172 | 0.104 | 0.019 | 0.5196 |
| 7 | 0.0515 | 0.9761 | 0.0172 | 0.1047 | 0.0191 | 0.5112 |
| 8 | 0.0528 | 0.9748 | 0.0171 | 0.1071 | 0.019 | 0.515 |
| 9 | 0.0527 | 0.9749 | 0.0171 | 0.1128 | 0.0191 | 0.5101 |
| 10 | 0.0539 | 0.9737 | 0.017 | 0.1196 | 0.0191 | 0.51 |
| 11 | 0.0548 | 0.9729 | 0.0169 | 0.1303 | 0.0192 | 0.5062 |
| 12 | 0.0596 | 0.968 | 0.0169 | 0.1279 | 0.0193 | 0.5011 |
| 13 | 0.0613 | 0.9661 | 0.0169 | 0.1368 | 0.0194 | 0.4956 |
| 14 | 0.0661 | 0.9606 | 0.017 | 0.1237 | 0.0194 | 0.4984 |
| 15 | 0.0676 | 0.9587 | 0.0168 | 0.1375 | 0.0195 | 0.4911 |
| 16 | 0.0698 | 0.956 | 0.0168 | 0.1434 | 0.0196 | 0.4867 |
| 17 | 0.0705 | 0.9552 | 0.017 | 0.1263 | 0.0194 | 0.4972 |
| 18 | 0.0752 | 0.949 | 0.0167 | 0.151 | 0.0195 | 0.4915 |
| 19 | 0.0781 | 0.9449 | 0.0168 | 0.1385 | 0.0195 | 0.4896 |
| 20 | 0.0827 | 0.9383 | 0.0167 | 0.1529 | 0.0196 | 0.4843 |
| 21 | 0.0822 | 0.939 | 0.0168 | 0.1407 | 0.0194 | 0.496 |
| 22 | 0.0827 | 0.9382 | 0.0169 | 0.1312 | 0.0196 | 0.4878 |
| 23 | 0.0923 | 0.9231 | 0.0167 | 0.1475 | 0.0195 | 0.4911 |
| 24 | 0.0907 | 0.9257 | 0.0169 | 0.131 | 0.0196 | 0.4864 |
| 25 | 0.0919 | 0.9237 | 0.017 | 0.1195 | 0.0195 | 0.4919 |
| 26 | 0.09 | 0.9268 | 0.0167 | 0.1551 | 0.0196 | 0.4862 |
| 27 | 0.0917 | 0.9241 | 0.0168 | 0.137 | 0.0195 | 0.4901 |
| 28 | 0.0977 | 0.9138 | 0.0169 | 0.128 | 0.0196 | 0.4838 |
| 29 | 0.0989 | 0.9116 | 0.0168 | 0.1372 | 0.0197 | 0.4826 |

| | | | | | | |
|---|---|---|---|---|---|---|
| 30 | 0.1007 | 0.9084 | 0.0524 | 0.8237 | 0.0528 | 0.8205 |
| 31 | 0.0774 | 0.5952 | 0.0501 | 0.8303 | 0.0507 | 0.8259 |
| 32 | 0.0743 | 0.6082 | 0.0479 | 0.8373 | 0.0487 | 0.8313 |
| 33 | 0.0727 | 0.6092 | 0.0479 | 0.8302 | 0.0488 | 0.8242 |
| 34 | 0.0724 | 0.6142 | 0.0482 | 0.8293 | 0.0485 | 0.8266 |
| 35 | 0.0683 | 0.6208 | 0.0464 | 0.8254 | 0.0469 | 0.8213 |
| 36 | 0.0679 | 0.6195 | 0.0462 | 0.8236 | 0.0472 | 0.8164 |
